# Adipocyte REVERBα dictates adipose tissue expansion during obesity

**DOI:** 10.1101/2020.09.23.308155

**Authors:** A. Louise Hunter, Charlotte E. Pelekanou, Nichola J. Barron, Rebecca C. Northeast, Antony Adamson, Polly Downton, Thomas Cornfield, Peter S. Cunningham, Leanne Hodson, Andrew S.I. Loudon, Richard D. Unwin, Mudassar Iqbal, David W. Ray, David A. Bechtold

## Abstract

The circadian clock component REVERBα is considered a dominant regulator of lipid metabolism, with global *Reverbα* deletion driving dysregulation of white adipose tissue (WAT) lipogenesis and obesity. However, a similar phenotype is not observed under adipocyte-selective deletion (*Reverbα^Flox2-6^Adipo^Cre^*), and transcriptional profiling demonstrates that, under basal conditions, direct targets of REVERBα regulation are limited, and include the circadian clock and collagen dynamics. Under high-fat diet (HFD) feeding, *Reverbα^Flox2-6^Adipo^Cre^* mice do manifest profound obesity, yet without the accompanying WAT inflammation and fibrosis exhibited by controls. Integration of the WAT REVERBα cistrome with differential gene expression reveals broad control of metabolic processes by REVERBα which is unmasked in the obese state. Adipocyte REVERBα does not drive an anticipatory daily rhythm in WAT lipogenesis, but rather modulates WAT activity in response to alterations in metabolic state. Importantly, REVERBα action in adipocytes is critical to the development of obesity-related WAT pathology and insulin resistance.

## INTRODUCTION

The mammalian circadian clock directs rhythms in behaviour and physiology to coordinate our biology with predictable changes in food availability and daily alternations between fasted and fed states. In this way, profound cycles in nutrient availability and internal energy state can be managed across multiple organ systems. A central circadian clock in the suprachiasmatic nuclei (SCN) drives daily rhythms in our behaviour (e.g. sleep/wake cycles) and physiology (e.g. body temperature), and orchestrates rhythmic processes in tissue systems across the body (Dibner et al., 2010; West and Bechtold, 2015). The molecular clock mechanism is also present in most cells and tissue. Normal peripheral tissue function and expression of a ‘complete’ rhythmic transcriptome requires local tissue clock activity, as well as input from the central clock and rhythmic systemic signals (Guo et al., 2005; Hughes et al., 2012; Kornmann et al., 2007; Koronowski et al., 2019; Lamia et al., 2008). The relative importance of each of these factors remains ill-defined. Nevertheless, it is clear that our rhythmic physiology and metabolic status reflects the interaction of clocks across the brain and body (West and Bechtold, 2015). Disturbance of this interaction, as occurs with shift work and irregular eating patterns, is increasingly recognised as a risk factor for metabolic disease and obesity (Broussard and Cauter, 2016; Kim et al., 2019).

Extensive work over the past 20 years has demonstrated that circadian clock function and its component factors are closely tied into energy metabolism (Bass and Takahashi, 2010; Reinke and Asher, 2019), with strong rhythmicity evident in cellular and systemic metabolic processes. Clock-metabolic coupling in peripheral tissues is adaptable, as demonstrated by classical food-entrainment studies (Damiola et al., 2000; Mistlberger, 1994), and by recent work showing that systemic perturbations such as cancer and high-fat diet feeding can re-programme circadian control over liver metabolism (Dyar et al., 2018; Masri et al., 2016). Plasticity therefore exists within the system, and the role of the clock in tissue and systemic responses to acute and chronic metabolic perturbation remains a critical question. The nuclear receptor REVERBα (NR1D1) is a core clock component, and has been highlighted as a key link between the clock and metabolism. REVERBα is a constitutive repressor, with peak expression in the latter half of the inactive phase (daytime in the nocturnal animal). In liver, REVERBα exerts repressive control over programmes of lipogenesis by recruiting the NCOR/HDAC3 co-repressor complex to metabolic target genes, such that loss of REVERBα or HDAC3 results in hepatosteatosis (Feng et al., 2011; Zhang et al., 2016, 2015). The selective functions of REVERBα in white adipose tissue (WAT) are not well-established and remain poorly understood. Early studies implicated an essential role of *Reverba* in adipocyte differentiation (Chawla and Lazar, 1993; Kumar et al., 2010); however, these findings are difficult to align with *in vivo* evidence. Indeed, pronounced adiposity and adipocyte hypertrophy are evident in *Reverbα*^-/-^ mice, even under normal feeding conditions (Delezie et al., 2012; Hand et al., 2015; Zhang et al., 2015). Moreover, daily administration of REVERBα agonists has been shown to reduce fat mass and WAT lipogenic gene expression in mice (Solt et al., 2012), despite concerns about off-targets actions of these agents (Dierickx et al., 2019). Given the links between circadian disruption and obesity, and the potential of REVERBα as a pharmacological target, we now define the role of REVERBα in dictating WAT metabolism.

Transcriptomic and proteomic profiling of WAT in global *Reverbα^-/-^* mice revealed an expected de-repression of lipid synthesis and storage programmes. However, in contrast, selective deletion of *Reverbα* in adipocytes did not result in dysregulation of WAT metabolic pathways. Loss of REVERBα activity in WAT did, however, significantly enhance adipose tissue expansion in response to HFD feeding; yet despite the exaggerated obesity, adipocyte-specific knockout mice were spared anticipated obesity-related pathology. Integration of transcriptomic data with the WAT REVERBα cistrome demonstrates that, under basal conditions, REVERBα activity is limited to a small set of direct target genes (enriched for extracellular matrix processes). However, REVERBα-regulatory control broadens to include lipid and mitochondrial metabolism pathways under conditions of obesity. Our data recast the role of REVERBα as a regulator responsive to the metabolic state of the tissue, rather than one which delivers an anticipatory daily oscillation to the WAT metabolic programme.

## RESULTS

### Adiposity and up-regulation of WAT lipogenic pathways in *Reverbα^-/-^* mice

We first examined the body composition of age-matched *Reverbα* global knockout (KO) (*Reverbα^-/-^*) mice and littermate controls (WT). In keeping with previous reports (Delezie et al., 2012; Hand et al., 2015), *Reverbα^-/-^* mice are of similar body weight to littermate controls (**Figure 1A**), yet carry an increased proportion of fat mass (KO: 24.2 ±3.0% of body weight; WT: 10.8 ±1.4%; mean ±SEM, P<0.01 Student’s t-test, n=12-14/group) and display adipocyte hypertrophy (**Figure 1B**), even when maintained on a standard chow diet. Metabolic phenotyping demonstrated expected day-night differences in food intake, energy expenditure, activity, and body temperature in both KO and WT controls, although genotype differences in day/night activity and temperature levels suggest some dampening of rhythmicity in the *Reverbα^-/-^* mice (**Figure 1 Supplemental A-E**). However, this is unlikely to account for the increased adiposity in these animals, and a previous study did not report significant genotype differences in these parameters (Delezie et al., 2012). This favours instead an altered energy partitioning within these mice, with a clear bias towards storing energy as lipid.

**FIGURE 1.**
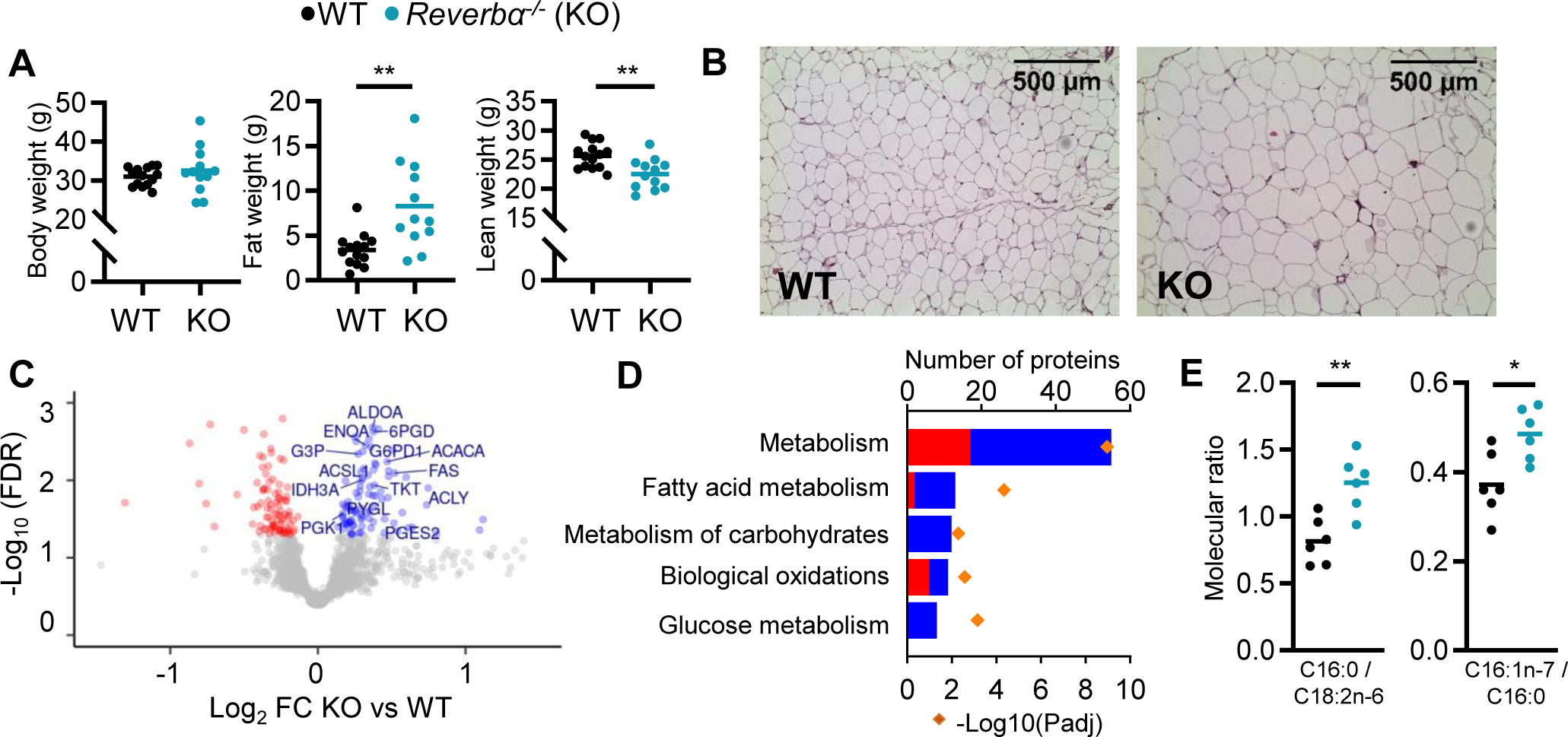
Global deletion of *Reverbα* results in obesity and increased adipose lipid synthesis. **A**. *Reverbα^-/-^* mice exhibit significantly greater fat mass relative to WT littermate controls. Body weight, fat mass and lean mass of 13-week old males (n=12-14/group). **B**. Increased fat mass in *Reverbα^-/-^* mice is reflected in adipocyte hypertrophy in gonadal white adipose tissue (gWAT) (representative x10 H&E images shown). **C**,**D**. gWAT from *Reverbα^-/-^* mice exhibits a programme of increased lipid synthesis. Proteomic profiling of gWAT depots (*Reverbα^-/-^* mice plotted relative to their respective weight-matched littermate controls, n=6/group (**C**)) shows deregulation of metabolic regulators and enrichment (**D**) of metabolic pathways (up- and down-regulated proteins shown in blue and red respectively). Top five (by protein count) significantly enriched Reactome terms shown. **E**. Analyses of fatty acid (FA) composition reveal increased *de novo* lipogenesis (reflected by C16:0/C18:2n ratio) and FA desaturation (reflected by C16:1n-7/C16:0 ratio) in gWAT of *Reverbα^-/-^* mice. n=6/group. Data presented as mean with individual data points (A, E). * P<0.05, **P<0.01, unpaired two-tailed t-test (A, E).

**FIGURE 1 SUPPLEMENTAL.**
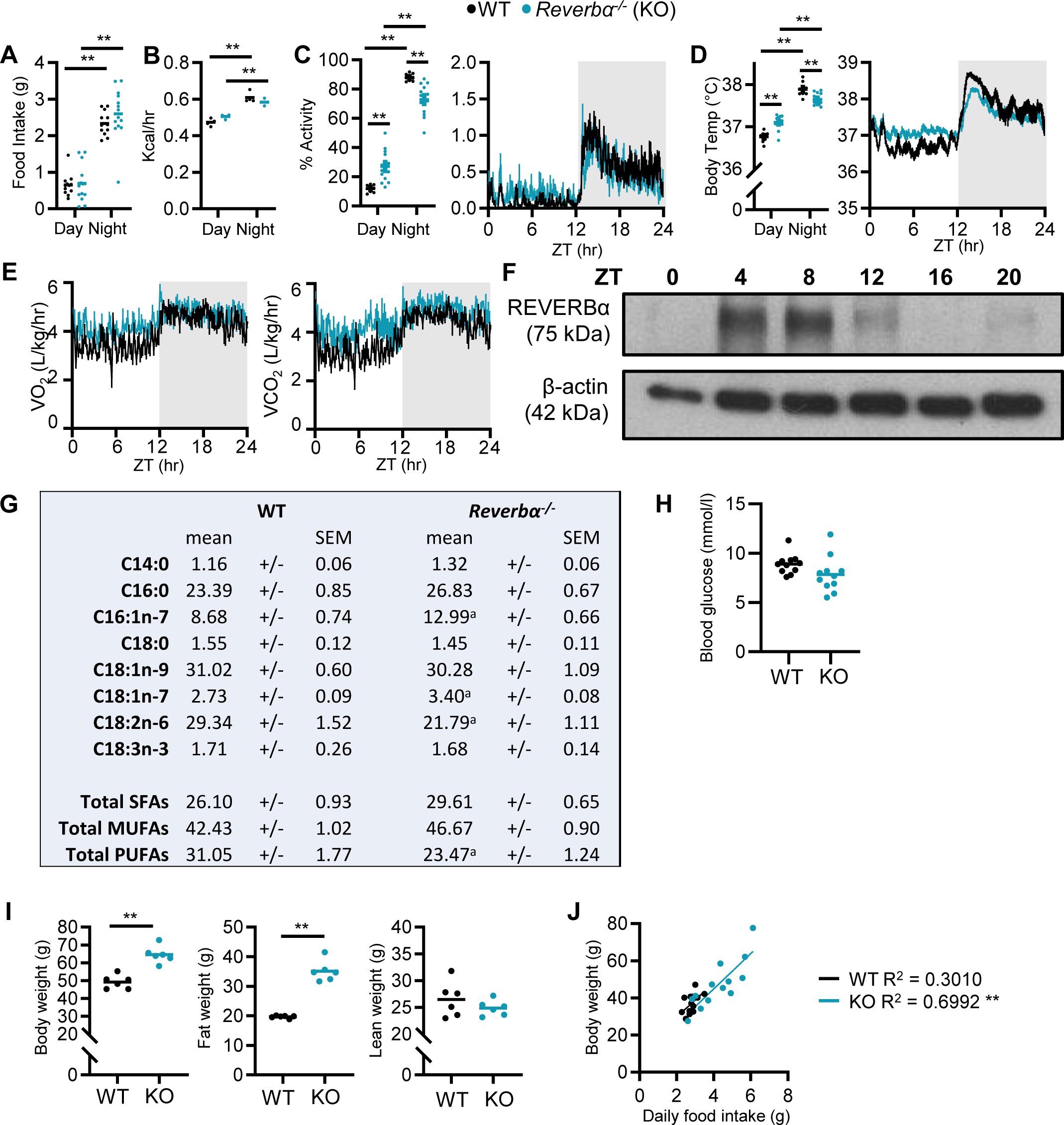
Rhythmic physiology and susceptibility to diet-induced obesity in *Reverbα^-/-^* mice. **A**-**E**. Under light:dark conditions, *Reverbα^-/-^* mice maintain robust diurnal rhythms in physiology and behaviour. Day/night food intake (**A**) (n=12-14/group), and energy expenditure (**B**) (n=3-4/group). Diurnal activity profile, mean activity (**C**) and body temperature (**D**) of adult male *Reverbα^-/-^* mice (activity is reported as the % daily activity, n=9-13/group). Diurnal profiles in oxygen consumption (VO_2_) and carbon dioxide production (VCO_2_) (**E**) (n=9-13/group). **F**. Western blot showing REVERBα expression in adipose tissue over 24 hours. **G**. Molar percentages of fatty acid species in WT and *Reverbα^-/-^* gWAT. n=6/group. ^a^P<0.05. **H**. Daytime (ZT6) fasted blood glucose levels (n=11/group). **I**. *Reverbα^-/-^* mice are highly susceptible to diet-induced obesity, showing significantly higher body weights and fat mass than control mice after 10 weeks of HFD feeding (n=6/group). **J**. In a separate study, food intake was tracked for individual mice over 3 weeks of HFD feeding (n = 13/group). Mean daily food intake in *Reverbα^-/-^* mice showed a significant positive correlation with body weight. Data presented as individual data points with mean (**A-D, H, I**), as mean +/- SEM (**F**), or as individual data points with line of best fit (**J**). **P<0.01, two-way ANOVA with Tukey’s multiple comparisons tests (**A**-**D**), unpaired t-tests with correction for multiple testing (**G**), unpaired two-tailed t-tests (**H**,**I**), linear regression (**J**).

To explore further the lipid storage phenotype, we undertook proteomic analysis of gonadal white adipose tissue (gWAT) collected at ZT8 (*zeitgeber* time, 8h after lights on), the time of normal peak in REVERBα expression in this tissue (**Figure 1 Supplemental F**). Isobaric tag (iTRAQ) labelled LC-MS/MS identified 2257 proteins, of which 182 demonstrated differential regulation (FDR<0.05) between WT and *Reverbα^-/-^* gWAT samples (n=6 weight-matched, 13-week old male mice/group) (**Figure 1C**). Differentially expressed proteins included influential metabolic enzymes, with up-regulation of metabolic processes detected on pathway enrichment analysis (**Figure 1D**). Importantly, and in line with the phenotype observed, increased NADPH regeneration (e.g. ME1, G6PDX), enhancement of glucose metabolism, (also likely reflecting increased glyceroneogenesis; e.g. PFKL, ALDOA), and up-regulation of fatty acid synthesis (e.g. ACYL, FAS, ACACA) all support a shunt towards synthesis and storage of fatty acids and triglyceride in the knockout mice. To validate this putative increase in local lipid synthesis, we quantified fatty acid species in gWAT, and indeed, the ratio of palmitic to linoleic acid (C16:0/C18:2n6), a marker of *de novo* lipogenesis, was significantly elevated in *Reverbα^-/-^* samples (**Figure 1E, Figure 1 Supplemental G**). Fatty acid profiling also revealed evidence of increased SCD1 activity (C16:1n-7/C16:0). Enhanced fatty acid synthesis in gWAT of mice lacking REVERBα may be in part driven by increased glucose availability and adipose tissue uptake as previously suggested (Delezie et al., 2012), although we do not observe elevated blood glucose levels in the *Reverbα^-/-^* animals (**Figure 1 Supplemental H**). The propensity to lipid storage is further highlighted by the substantial obesity, compared to littermate controls, displayed by *Reverbα^-/-^* mice when challenged with 10 weeks of high-fat diet (HFD) (**Figure 1 Supplemental I**; Delezie et al., 2012; Hand et al., 2015). Interestingly, we observed a strong positive correlation between body weight and daily intake of HFD in the *Reverbα^-/-^* mice (**Figure 1 Supplemental J**), suggesting that HFD-induced hyperphagia exacerbates weight gain and obesity in the *Reverbα^-/-^* mice.

### Limited impact of adipocyte-selective *Reverbα* deletion under basal conditions

To define the role of REVERBα specifically within WAT, we generated a new mouse line with loxP sites flanking *Reverbα* exons 2-6 (*Reverbα^Flox2-6^*), competent for Cre-mediated conditional deletion (**Figure 2A**; Hunter et al., 2020). We crossed this mouse with the well-established adiponectin Cre-driver line (*Adipo^Cre^*; Eguchi et al., 2011; Jeffery et al., 2014) to delete *Reverbα* selectively in adipocytes. This new line results in loss of *Reverbα* mRNA (**Figure 2B**) and protein (**Figure 2C**) expression in adipose tissue depots, as well as coordinate de-repression of *Bmal1,* upon Cre-mediated recombination.

**FIGURE 2.**
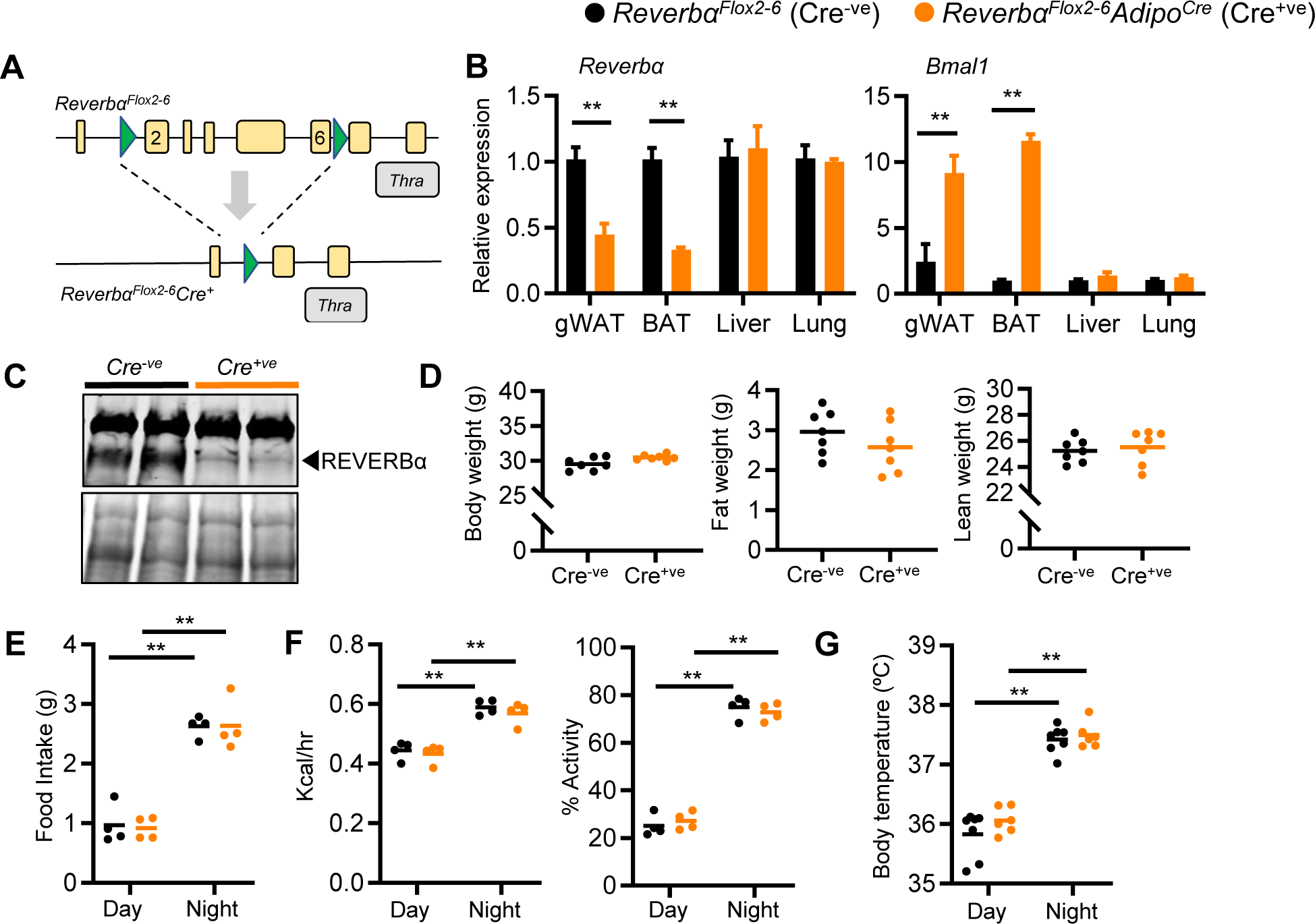
Impact of adipose *Reverbα* deletion is limited under normal conditions. **A**. Targeting strategy for LoxP site integration flanking exons 2-6 of the *Nr1d1* (*Reverbα*) locus. **B**. *Reverbα* and *Bmal1* gene expression in gWAT, brown adipose (BAT), liver and lung in *Reverbα^Flox2-6^* (Cre^-ve^) and *Reverbα^Flox2-6^Adipo^Cre^* (Cre^+ve^) mice (n=4-5/group). **C**. REVERBα protein expression (arrowhead) in Cre^-ve^ and targeted Cre^+ve^ mice. Lower blot shows Ponceau S protein staining. **D**. Body weight, fat mass and lean mass in 13-week old Cre^-ve^ and Cre^+ve^ male mice (n=7/group). **E**-**G**. Both *Reverbα^Flox2-6^Adipo^Cre^* Cre^+ve^ and Cre^-ve^ mice demonstrate diurnal rhythms in behaviour and physiology, with no genotype differences observed in food intake (**E**), energy expenditure and daily activity (**F**) or body temperature (**G**) in 13-week old males (n=4-7/group). Data presented as mean +/-SEM (**B**,) or as mean with individual data points (**D-G**). *P<0.05, **P<0.01, unpaired t-tests corrected for multiple comparisons (**B**), unpaired t-tests (**D**), two-way ANOVA with Tukey’s multiple comparisons tests (**E**-**G**).

**FIGURE 2 SUPPLEMENTAL.**
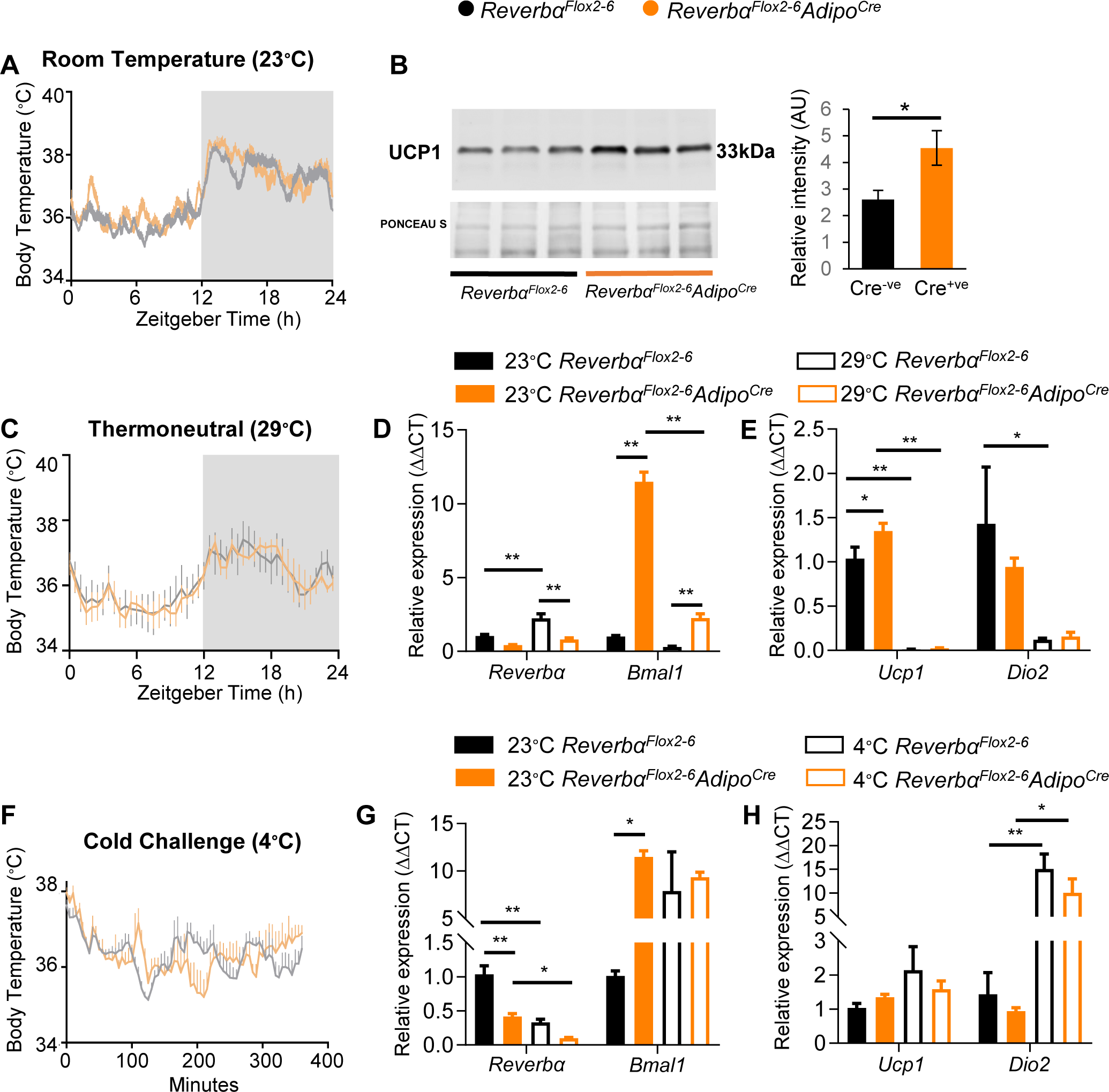
Loss of *Reverba* expression in brown adipocytes does not alter body temperature. **A-B**. Housing under standard laboratory conditions did not alter daily profiles in body temperature in the *Reverbα^Flox2-6^Adipo^Cre^* Cre^+ve^ mice, when compared to control Cre^-ve^ littermate controls (**A**; n=5-6/group), despite showing increased UCP1 expression (**B**; n=3/group). **C**. No intergenotype genotype differences were observed in body temperature profiles recorded from mice housed under thermoneutral conditions (28-30^∘^C) for 3 weeks (n=5-6/group). **D-E**. Brown adipose tissue (BAT) gene expression studies (qPCR) demonstrated expected de-repression of *Bmal1* expression in Cre^+ve^ mice, and expected reduction in *Ucp1* expression at thermoneutral conditions in both genotypes (compared to room temperature) (n=5-6/group). **F**. *Reverbα^Flox2-6^Adipo^Cre^* Cre^+ve^ mice and control littermates were exposed to an acute cold challenge (4^∘^C for 6 h) with body temperature recording throughout (n=5-6/group). No genotype difference in thermogenic response was observed. **G-H.** As previously reported, *Reverbα* expression was reduced by cold exposure; however, no genotype differences were observed cold-induced increases in *Ucp1* or *Dio2* gene expression (n=5-6/group). Data presented as mean +/-SEM. *P<0.05, **P<0.01, Student’s t-test (**B**), 2-way ANOVA with Tukey’s multiple comparisons tests (**D**, **E**, **G**, **H**).

In marked contrast to global *Reverbα^-/-^* mice, adult *Reverbα^Flox2-6^Adipo^Cre^* mice did not show an increase in adiposity when maintained on a standard chow diet (**Figure 2D**; n=7/group), with no differences in mean body weight, fat and lean mass observed. In parallel with this, we saw no differences in daily patterns of food intake, energy expenditure, activity levels or body temperature in matched *Reverbα^Flox2-6^Adipo^Cre^* and control (*Reverbα^Flox2-6^*) mice (**Figure 2D**,**E**).

As brown adipose tissue (BAT) makes an important contribution to whole body energy metabolism, we studied the thermoregulation of *Reverbα^Flox2-6^Adipo^Cre^* mice in greater detail. It has previously been proposed that REVERBα is key in conferring circadian control over thermogenesis, through its repression of uncoupling protein 1 (UCP1) (Gerhart-Hines et al., 2013). However, we saw no genotype differences in thermoregulation between *Reverbα^Flox2- 6^Adipo^Cre^* and *Reverbα^Flox2-6^* mice (**Figure 2 Supplemental A-E**). Despite increased BAT UCP1 expression, no differences in body temperature profiles were observed between Cre^-ve^ and Cre^+ve^ mice when housed under normal laboratory temperature (∼22°C) nor when placed under thermoneutral conditions (29°C) for >14days. Moreover, Cre^-ve^ and Cre^+ve^ mice did not differ in their thermogenic response to an acute cold challenge (4°C for 6hr) (**Figure 2 Supplemental F-H**). Therefore, the minimal impact on body composition of adipose-targeted *Reverbα* deletion cannot be explained by altered BAT thermogenic activity, and moreover, these data challenge existing theories about the role of *Reverbα* in thermoregulation.

### In normal WAT, REVERBα-regulated targets are limited to clock and collagen genes

To investigate adipocyte-specific *Reverbα* activity, we performed RNA-seq at ZT8 (n=6/group) in both *Reverbα^-/-^* and *Reverbα^Flox2-6^Adipo^Cre^* mouse lines. Global *Reverbα* deletion had a large effect on the gWAT transcriptome, with 4163 genes showing significant differential expression (FDR<0.05) between *Reverbα^-/-^* mice and age- and weight-matched WT littermate controls (**Figure 3A**). Pathway enrichment analysis demonstrated that these changes are dominated by metabolic genes (**Figure 3B(i)**), with lipid metabolism and the TCA cycle emerging as prominent processes (**Figure 3B(ii)**). Thus, the gWAT transcriptome in *Reverbα^-/-^* mice is concordant with the phenotype, and the gWAT proteome, in demonstrating up-regulation of lipid accumulation and storage processes.

**FIGURE 3.**
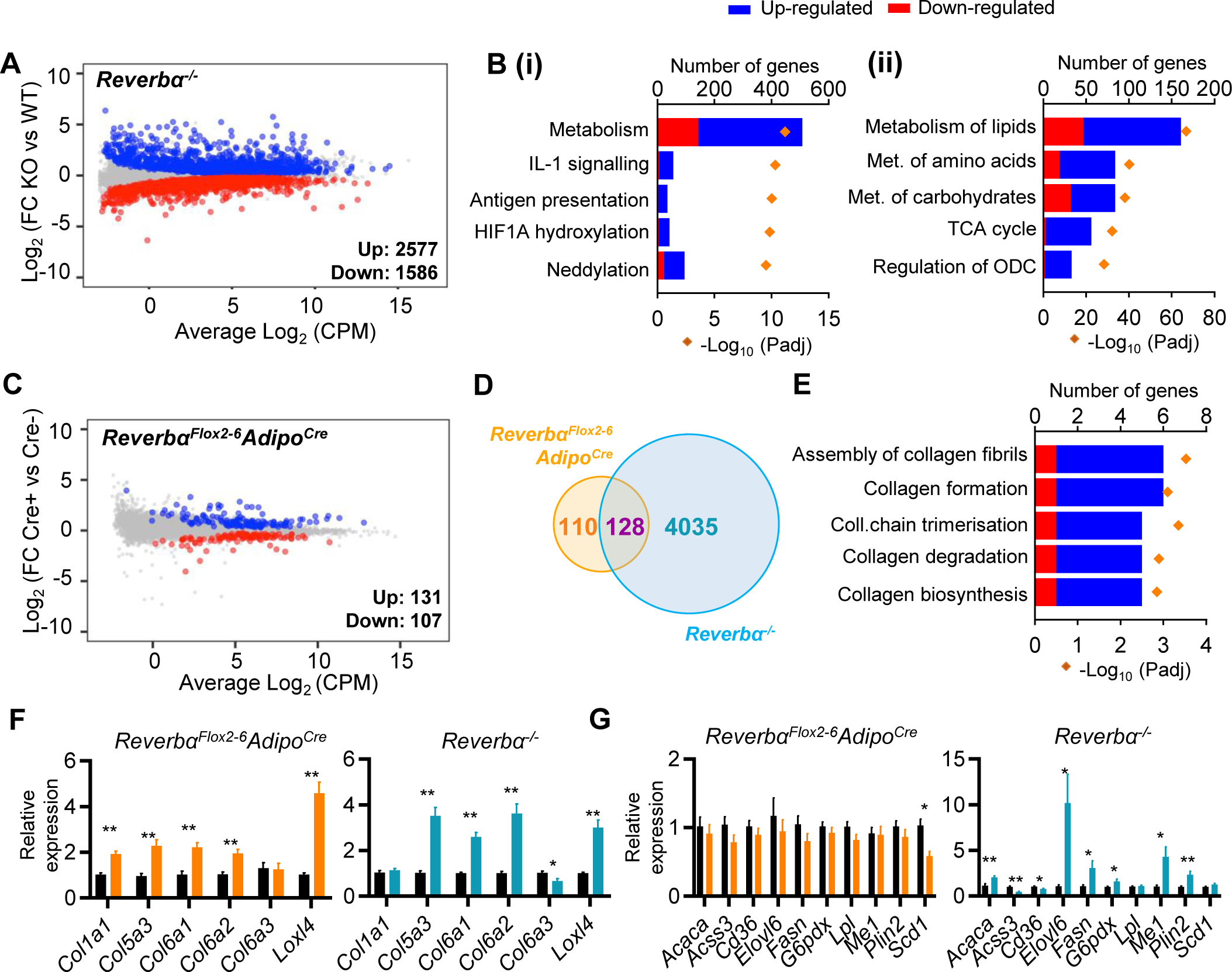
Global or adipose-specific *Reverbα* deletion produces distinctive gene expression profiles. **A, B**. *Reverbα^-/-^* gWAT demonstrates extensive remodelling of the transcriptome and up-regulation of metabolic pathways. Mean-difference (MD) plot (**A**) showing significantly (FDR<0.05) up- (blue) and down- (red) regulated genes in gWAT of *Reverbα^-/-^* mice compared to littermate controls (n=6/group). Pathway analysis (**B**) of significantly differentially expressed genes (FDR<0.05): top five (by gene count) significantly enriched Reactome terms shown (**B(i)**), top 5 metabolic pathways shown (**B(ii)**). Up-regulated genes in blue, down- regulated in red. ODC = ornithine decarboxylase. **C**. By contrast, RNA-seq demonstrates modest remodelling of the transcriptome in gWAT of *Reverbα^Flox2-6^Adipo^Cre^* mice. MD plot, n=6/group. **D**. Venn diagram showing overlap of differentially-expressed (DE) genes in *Reverbα^Flox2-6^Adipo^Cre^* and *Reverbα^-/-^* gWAT. **E**. Pathway analysis of 128 commonly DE genes. Top five (by gene count) significantly enriched Reactome terms shown. **F**,**G**. Collagen genes are commonly up-regulated in both genotypes (**F**), whilst genes of lipid metabolism are not DE in *Reverbα^Flox2-6^Adipo^Cre^* (**G**). gWAT qPCR, n=6-7/group. Data presented as mean +/- SEM (**F, G**). *P<0.05, **P<0.01, unpaired t-tests corrected for multiple comparisons (**F, G**).

**FIGURE 3 SUPPLEMENTAL.**
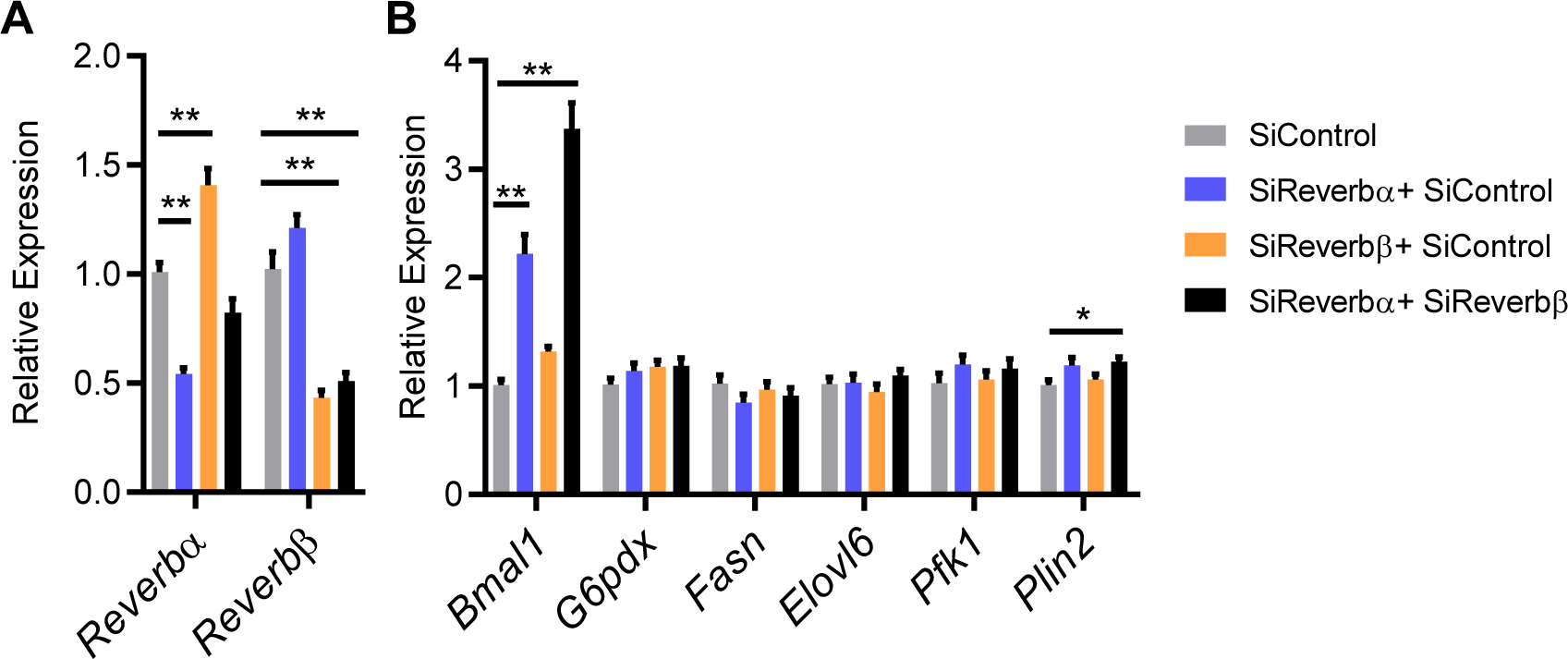
Impact of *Reverbα* and *Reverbβ* loss *in vitro*. **A, B.** Double knockdown of *Reverbα* and *Reverbβ* in differentiated 3T3-L1 cells (**A**) results in marked de- repression of *Bmal1* expression but minimal effects on expression of typically pro-lipogenic genes (**B**) (data compiled from three replicated knockdown experiments, n=8-9/treatment group). Data presented as mean +/- SEM. *P<0.05, **P<0.01, one-way ANOVA with Dunnett’s multiple comparisons tests (**A**, **B**).

By contrast, and consistent with the absence of an overt phenotype, only a small genotype effect on the transcriptome was observed when comparing gWAT RNA-seq from *Reverbα^Flox2- 6^Adipo^Cre^* and *Reverbα^Flox2-6^* littermates (**Figure 3C**; n=6/group). Here, 238 genes showed significant differential expression between genotypes, of which 128 were also differentially regulated in the WAT analysis of global *Reverbα^-/-^* mice (**Figure 3D**). These 128 common genes included circadian clock components (*Bmal1, Clock, Cry2, Nfil3*), whilst pathway analysis also revealed collagen formation/biosynthesis processes to be significantly enriched (**Figure 3C**). Regulation of the molecular clock is expected, but the discovery of collagen dynamics as a target of REVERBα regulatory action in adipocytes has not been previously recognised. We validated consistent up-regulation of collagen and collagen-modifying genes in *Reverbα^-/-^* and *Reverbα^Flox2-6^Adipo^Cre^* gWAT by qPCR (**Figure 3F**).

It is notable that *Reverbα^Flox2-6^Adipo^Cre^* mouse gWAT exhibits neither enrichment of lipid metabolic pathways, nor de-regulation of individual key lipogenic genes, previously identified as REVERBα targets (**Figure 3E,G**) (Feng et al., 2011; Zhang et al., 2016, 2015). These findings suggest lipogenic gene regulation may be a response to system-wide changes in energy metabolism in the *Reverbα^-/-^* animals, and challenge current understanding of REVERBα action.

Work in liver has suggested that the REVERBα paralogue, REVERBβ, contributes to the suppression of lipogenesis, and that concurrent REVERBβ deletion amplifies the impact of REVERBα loss (Bugge et al., 2012). We therefore performed double knock-down of *Reverbα* and *Reverbβ* in differentiated 3T3-L1 cells (**Figure 3 Supplemental A**). Whilst double knock- down produced greater *Bmal1* de-repression than either *Reverbα* or *Reverbβ* knock-down alone, it did not lead to de-repression of lipogenic genes previously considered REVERBα targets (**Figure 3 Supplemental B**). This suggests that compensation by *Reverbβ* does not underlie the mild phenotype observed in the *Reverbα^Flox2-6^Adipo^Cre^* mice.

Thus, whilst global REVERBα targeting produces an adiposity phenotype with up-regulation of WAT lipogenesis and lipid storage, this is not seen when REVERBα is selectively targeted in adipose alone. The distinction is not due to loss of *Reverbα* expression in brown adipose, and is not due to compensatory REVERBβ action. Taken together, our data suggest that under a basal metabolic state, the adipose transcriptional targets under direct REVERBα control are in fact limited to core clock function and collagen dynamics. REVERBα is not a major repressor of lipid metabolism in this setting. This also suggests that the enhanced lipid accumulation phenotype of *Reverbα^-/-^* adipose tissue is either independent from adipose REVERBα entirely, or that the action of REVERBα in adipose is context-dependent.

### Diet-induced obesity reveals a broader WAT phenotype in tissue-specific REVERBα deletion

Studies in liver tissue have demonstrated reprogramming of both nuclear receptor and circadian clock factor activity by metabolic challenge (Eckel-Mahan et al., 2013; Goldstein et al., 2017; Guan et al., 2018; Quagliarini et al., 2019). Both our data here, and previous reports (Delezie et al., 2012; Feng et al., 2011; Hand et al., 2015; Le Martelot et al., 2009; Preitner et al., 2002), highlight that the normal chow-fed *Reverbα^-/-^* mouse is metabolically abnormal. The emergence of the collagen dynamics as a direct REVERBα target and exaggerated diet-induced obesity evident in *Reverbα^-/-^* mice supports a role for REVERBα in regulating adipose tissue expansion under obesogenic conditions. To test this, *Reverbα^Flox2-6^Adipo^Cre^* and *Reverbα^Flox2-6^* mice were provided with HFD for 16 weeks to drive obesity and WAT expansion. Indeed, compared to their controls, *Reverbα^Flox2-6^Adipo^Cre^* mice exhibited greater weight gain and adiposity in response to HFD feeding (**Figure 4A,B**). Of note, divergence between control and *Reverbα^Flox2-6^Adipo^Cre^* mice became clear only after long-term HFD-feeding (beyond ∼13 weeks), a time at which body weight gain plateaus in control mice. This contrasts substantially with *Reverbα^-/-^* mice, which show rapid and profound weight gain from the start of HFD feeding (Hand et al., 2015). The stark difference in progression and severity of diet-induced obesity is likely due (at least in part) to the HFD-induced hyperphagia, which is observed in *Reverbα^-/-^* mice (WT food intake 2.92 ±0.10g HFD/day/mouse; KO 3.74 ±0.21g, P=0.0014, Student’s T- test, n=21/genotype), but not in *Reverbα^Flox2-6^Adipo^Cre^* mice (Cre^-ve^ 2.99 ±0.61g HFD/day/mouse; Cre^+ve^ 3.01 ±0.60g, P>0.05, n=8/genotype). Nevertheless, both models highlight that loss of REVERBa increases capacity for increased lipid storage and adipose tissue expansion under obesogenic conditions.

**FIGURE 4.**
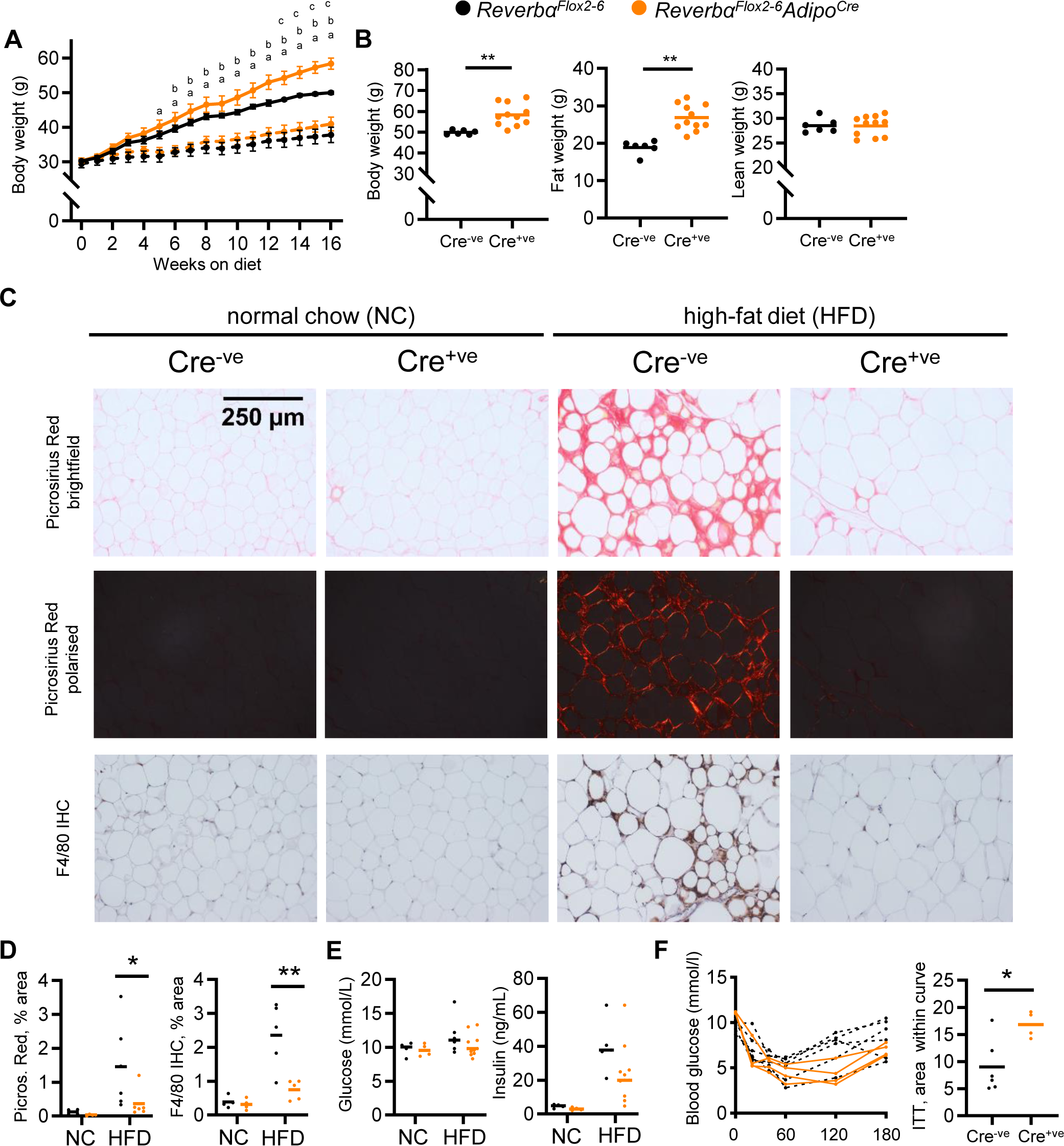
Diet-induced obesity unmasks a role for REVERBα in the regulation of adipose expansion. **A**, **B**. High-fat diet leads to exaggerated adiposity in *Reverbα^Flox2-6^Adipo^Cre^* mice. Body weight track of Cre^-ve^ and Cre^+ve^ male mice on high-fat diet (solid line) or normal chow (dashed line) (**A**) (^a^P<0.05: Cre^+ve^ NC vs HFD; ^b^P<0.05, Cre^-ve^ NC vs HFD; ^c^P<0.05, Cre^+ve^ HFD vs Cre^-ve^ HFD); total body, fat and lean weight after 16 weeks in the high-fat diet group (**B**). **C**,**D**. On histological examination of WAT, HFD-fed Cre^+ve^ mice display less fibrosis and inflammation than Cre^-ve^ littermates. Representative Picrosirius Red and F4/80 immunohistochemistry images (x20 magnification) (**C**), quantification of staining across groups, each data point represents the mean value for each individual animal (**D**). **E**,**F**. Despite increased adiposity, HFD-fed Cre^+ve^ mice display greater insulin sensitivity than Cre^-ve^ controls. Terminal blood glucose and insulin levels (animals culled 2hrs after food withdrawal) in NC and HFD-fed in *Reverbα^Flox2-6^Adipo^Cre^* Cre^-ve^ (black) and Cre^-ve^ (orange) mice (**E**). Blood glucose values for individual animals and area within curve for 16-week HFD-fed *Reverbα^Flox2- 6^Adipo^Cre^* Cre^-ve^ and Cre^-ve^ mice undergoing insulin tolerance testing (ITT) (**F**). Data presented as mean +/- SEM (**A**) or as individual data points with mean (**B**, **D**, **E**, **F**). *P<0.05, **P<0.01, two-way repeated measures ANOVA with Tukey’s multiple comparisons tests (**A**), two-way ANOVA with Sidak’s multiple comparisons tests (**D**, **E**), unpaired two-tailed t-test (**B**, **F**). n=4-11/group for all panels.

**FIGURE 4 SUPPLEMENTAL.**
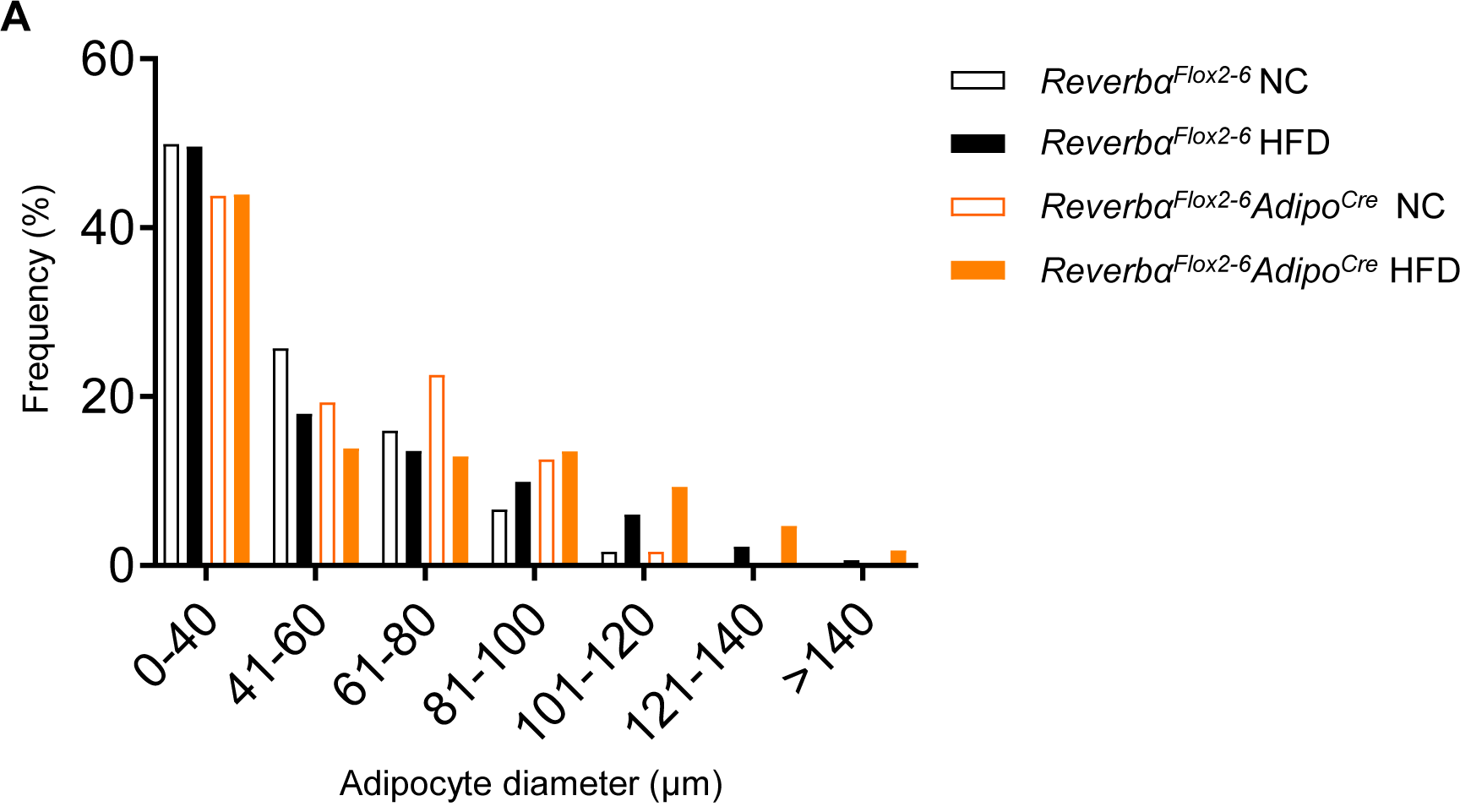
Adipocyte size in *Reverbα^Flox2-6^* and *Reverbα^Flox2-6^Adipo^Cre^* mice. **A**. Quantification of WAT adipocyte size in NC- and HFD-fed *Reverbα^Flox2-6^* and *Reverbα^Flox2-6^Adipo^Cre^* mice demonstrates no between-genotype differences in size distribution. n=4-6/group. Data presented as mean. Two-way ANOVA with Tukey’s multiple comparisons tests.

Despite the enhanced diet-induced obesity, HFD-fed *Reverbα^Flox2-6^Adipo^Cre^* mice showed little evidence of typical obesity-related pathology. Histological assessment of gWAT after 16- weeks of HFD feeding revealed widespread adipose tissue fibrosis (Picrosirius Red staining of collagen deposition under normal and polarised light) and macrophage infiltration (F4/80 immunohistochemistry) in obese control mice, but these features were not seen in *Reverbα^Flox2-6^Adipo^Cre^* mice (**Figure 4C,D**). Furthermore, we saw evidence of preserved insulin sensitivity in the HFD-fed *Reverbα^Flox2-6^Adipo^Cre^* mice, with neither circulating glucose nor insulin being higher than *Reverbα^Flox2-6^* littermate controls (**Figure 4E**), despite carrying significantly greater fat mass. Indeed, on insulin tolerance testing, HFD-fed *Reverbα^Flox2- 6^Adipo^Cre^* mice demonstrated more marked hypoglycaemia than HFD-fed controls (**Figure 4F**). We saw no differences in adipocyte size between the two genotypes, indicating that our observations did not simply reflect greater adipocyte hyperplasia in the *Reverbα^Flox2-6^Adipo^Cre^* mice (**Figure 4 Supplemental A**).

Therefore, under long-term HFD-feeding conditions, adipose-targeted *Reverbα* deletion results in continued adipose tissue expansion accompanied by a healthier metabolic phenotype with reduced adipose inflammation and fibrosis, and preserved systemic insulin sensitivity. Importantly, these findings also suggest that the regulatory influence of REVERBα is context-dependent, with the metabolic impact of adipose-targeted *Reverbα* deletion revealed by transition to an obese state.

### REVERBα-dependent gene regulation is reprogrammed by obesity

We next performed RNA-seq on gWAT collected at ZT8 from *Reverbα^Flox2-6^Adipo^Cre^* and *Reverbα^Flox2-6^* littermate controls fed either normal chow (NC) or HFD for 16 weeks (NC, n=4/group; HFD, n=6/group). As expected, HFD-feeding had a substantial impact on the gWAT transcriptome in both Cre^-ve^ and Cre^+ve^ animals (i.e. NC vs HFD comparison within each genotype; **Figure 5A**). Under NC feeding conditions, we again observed only a small genotype effect on the transcriptome, and as before, differentially-expressed genes included core clock genes (*Bmal1, Nfil3, Npas2, Clock*) and those of collagen synthesis pathways (**Figure 5B**). However, obesity revealed a substantial genotype effect, with 3061 genes differentially expressed (1706 up, 1355 down) in HFD-fed *Reverbα^Flox2-6^Adipo^Cre^* mice versus HFD-fed *Reverbα^Flox2-6^* controls (**Figure 5B**), and 1704 genes showing a significant (α<0.05) diet-genotype interaction (stageR specific interaction analysis; Van den Berge et al., 2017). Of these 1704 genes, those up-regulated in obese *Reverbα*-deficient adipose were strongly enriched for metabolic pathways, whilst down-regulated genes showed weak enrichment of ECM organisation processes (**Figure 5C**). To examine how loss of *Reverbα* alters adipose tissue response to diet-induced obesity, we compared directly those processes which showed significant obesity-related dysregulation in control mice (**Figure 5D**). While HFD-feeding caused a profound down-regulation (vs NC conditions) of metabolic pathways in the WAT of control mice, this was not observed in *Reverbα^Flox2-6^Adipo^Cre^* mice. By contrast, HFD-feeding led to an up-regulation of immune pathways in both genotypes (**Figure 5D**). Thus, transcriptomic profiling correlates with phenotype in suggesting that WAT function and metabolic activity is protected from obesity-related dysfunction in the *Reverbα^Flox2-6^Adipo^Cre^* mice, and that the impact of adipose *Reverbα* deletion is dependent on system-wide metabolic state.

**FIGURE 5.**
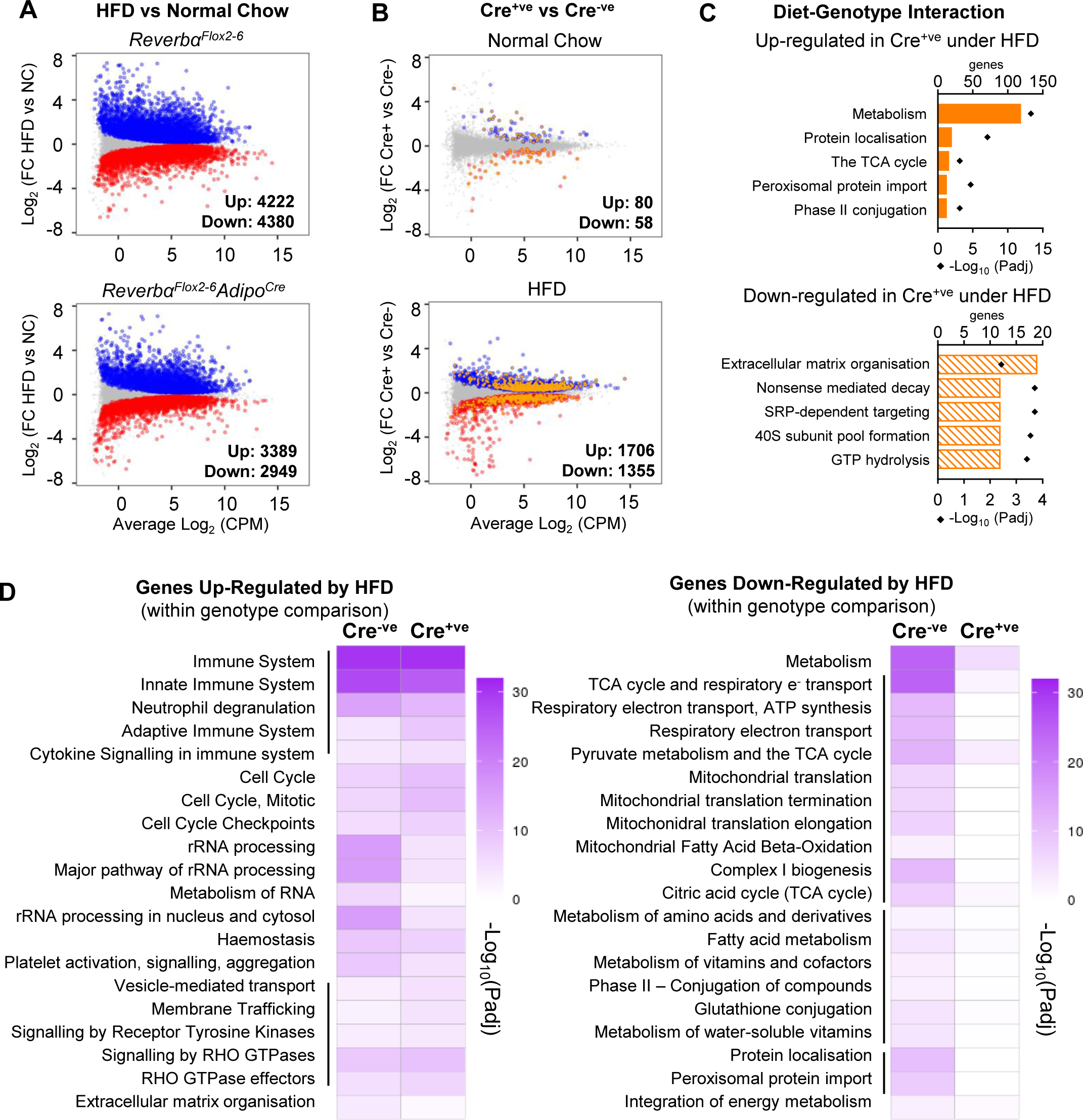
Under conditions of obesity, REVERBα repression extends to metabolic pathways. **A**. High-fat diet dramatically remodels the WAT transcriptome. RNA-seq (n=4-6/group) was performed in gWAT from Cre^-ve^ and Cre^+ve^ male mice fed normal chow (NC) or high-fat diet (HFD) for 16 weeks. MD plots show genes significantly (FDR<0.05) up-regulated (blue) or down-regulated (red) by HFD in each genotype. **B**. With HFD, the REVERBα-responsive gWAT transcriptome broadens. MD plots show effect of genotype in NC (top panel) and HFD (lower panel) feeding conditions. Genes where stageR detects a significant (α=0.05) genotype- diet interaction highlighted in orange. **C**. Reactome pathway analysis of genes up- or down-regulated in Cre^+ve^ gWAT under HFD conditions, where this diet-genotype interaction is also detected. Top five (by gene count) significantly enriched terms shown. **D**. Adipose-targeted deletion of *Reverbα* attenuates the normal HFD- induced down-regulation of metabolic pathways. Heatmaps show enrichment (-log_10_(Padj)) of Reactome pathways in genes up-regulated (left) or down-regulated (right) by HFD feeding in Cre^-ve^ and Cre^+ve^ gWAT. Top twenty (by gene count in Cre^-ve^ group) significantly enriched terms shown.

### Integration of differential gene expression with the WAT cistrome reveals state- dependent regulation of metabolic targets by REVERBα

To examine the mechanism of REVERBα regulation of the WAT metabolic programme in HFD- fed mice, we analysed the relationship between differentially expressed genes revealed by our RNA-seq studies and the WAT REVERBα cistrome. We identified primary REVERBα target genes by comparing genes changing in *Reverbα^Flox2-6^Adipo^Cre^* gWAT (relative to control mice, under both NC and HFD conditions) with genes changing in *Reverbα^-/-^* gWAT. To define the cistrome, we used raw published gWAT ChIP-seq data (Zhang et al., 2015) to call 2,354 high-stringency REVERBα peaks. To infer which genes might be direct targets of REVERBα repression, we employed a custom Python script that calculates the enrichment of differentially expressed gene sets in spatial relation to identified transcription factor binding sites (**Figure 6 Supplemental A**) (Hunter et al., 2020; Yang et al., 2019), over all genes in the genome.

**FIGURE 6.**
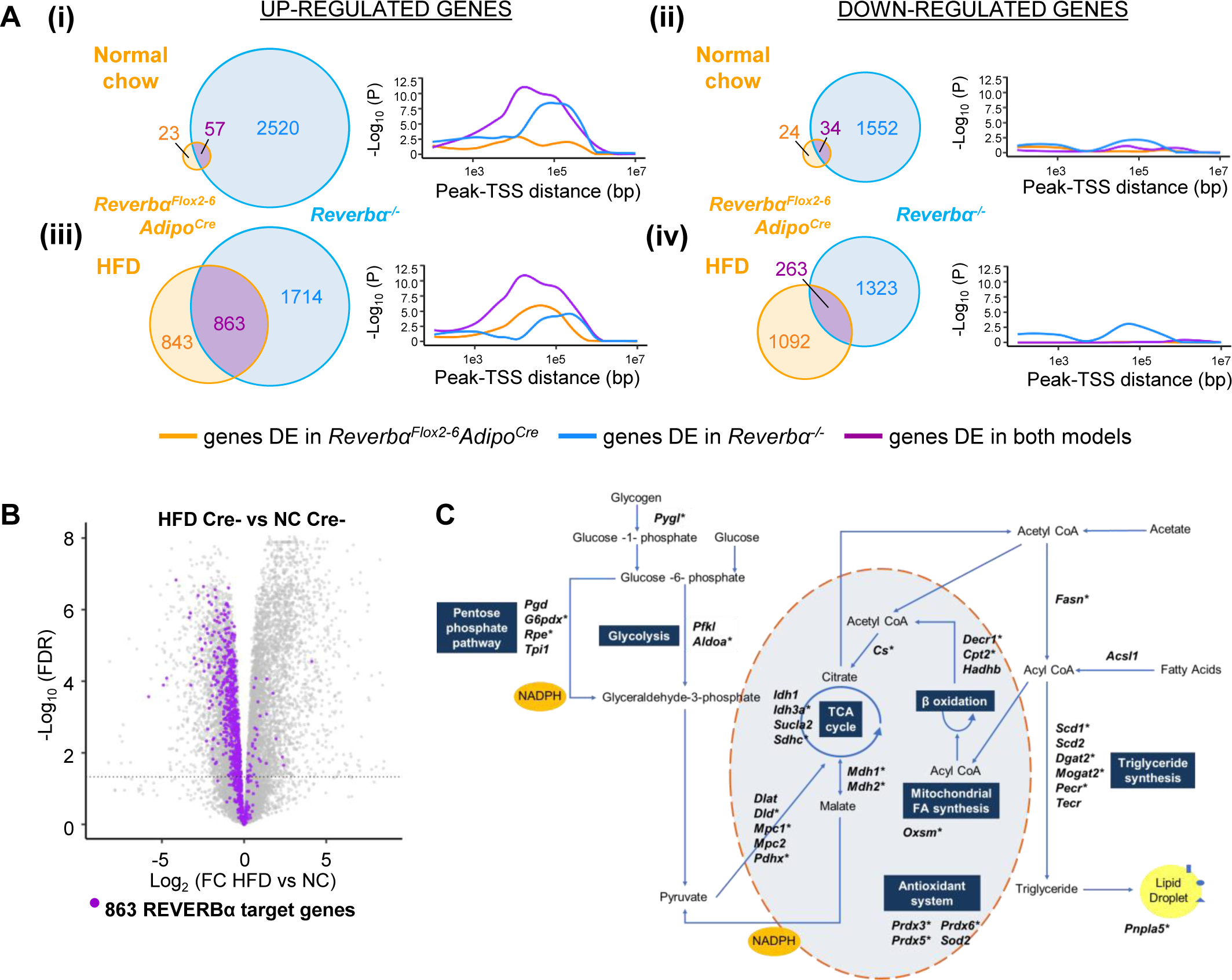
REVERBα-regulated targets, unmasked by obesity, associate with gWAT REVERBα binding sites. **A**. Genes commonly up-regulated by *Reverbα* loss, in both the basal and obese state, show the strongest association with gWAT REVERBα ChIP-seq peaks. Venn diagrams show overlap of differentially-expressed (DE) genes in *Reverbα^Flox2-6^Adipo^Cre^* Cre^+ve^ mice (under NC (**i, ii**) and HFD (**iii**,**iv**) conditions) with differentially- expressed genes in *Reverbα^-/-^* mice (each compared to their respective controls). Plots show –log_10_(P-value for enrichment) for each gene cluster at increasing distances from stringent REVERBα gWAT ChIP-seq peaks (FE>7.5 over input). See Figure 6 Supplemental (A) for schematic. DE genes separated by direction of regulation (up or down). **B**,**C**. REVERBα targets are also down-regulated in obesity, and include important lipid and mitochondrial metabolic regulators. Volcano plot (**B**) highlighting effect of HFD (in intact (Cre^-ve^) animals) of the 863 REVERBα target genes from **A(iii)**. Metabolic map illustrating REVERBα targets (**C**). Genes showing significantly down-regulation by HFD (in Cre^-ve^ animals) are starred*.

**FIGURE 6 SUPPLEMENTAL.**
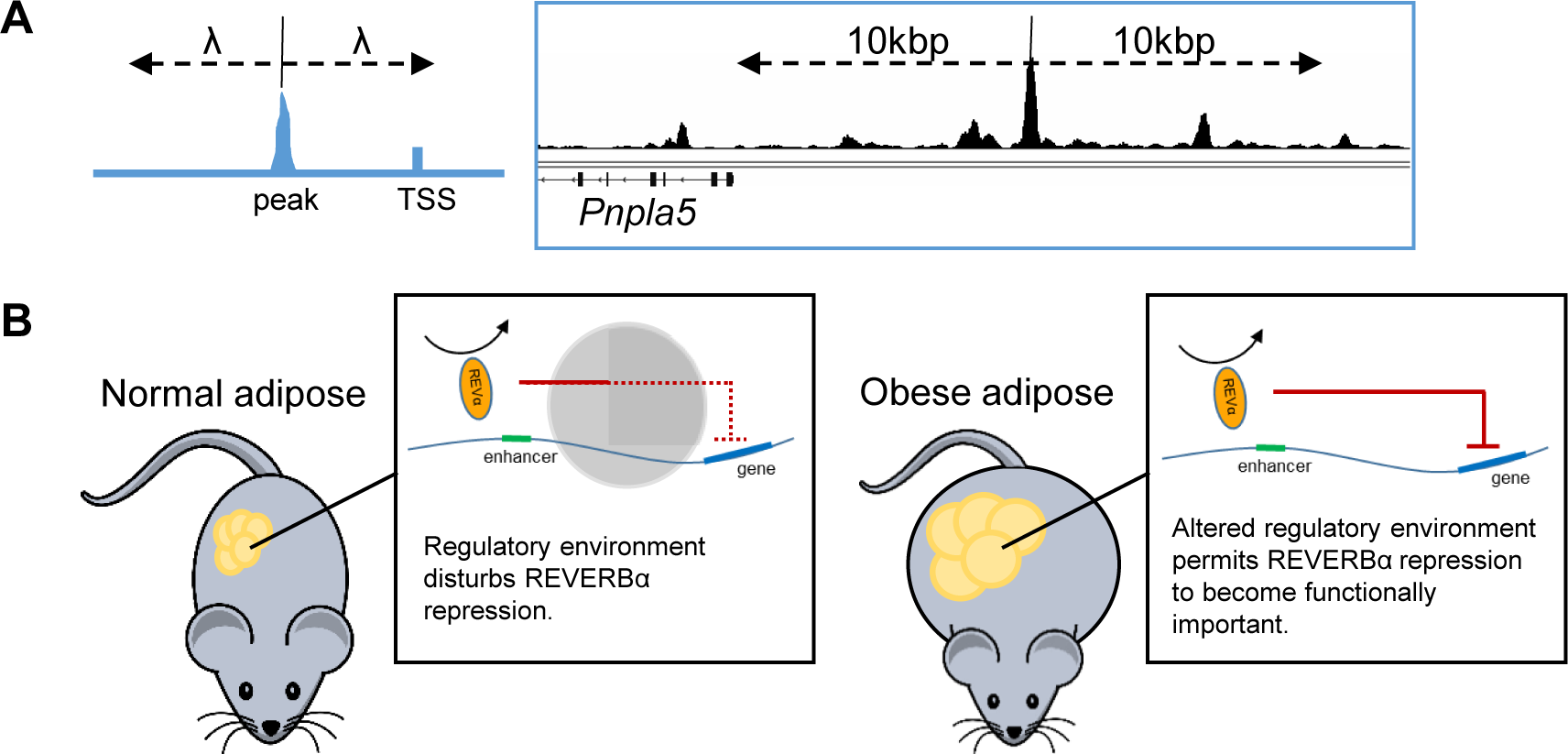
Relationship between REVERBα binding sites and gene regulation. **A**. Schematic of peaks-genes analysis. ChIP-seq peaks are extended by user-specified distances (λ) either side of the peak centre. Enrichment of the genes of interest, within all genes whose transcription start site (TSS) is detected within λ, is assessed by a hypergeometric test. The TSS of *Pnpla5*, for example, lies within 10kbp of a REVERBα ChIP-seq peak. **B**. Proposed model. In normal conditions, there is a large number of genes over which REVERBα repressive control is not apparent, likely because the regulatory environment (chromatin state, presence of other regulators) blocks this interaction or renders it redundant. In obese adipose, alterations to the regulatory environment (e.g. chromatin remodelling) are permissive to REVERBα activity.

Under NC conditions, only small sets of genes were up- or down-regulated in both *Reverbα^Flox2-6^Adipo^Cre^* and *Reverbα^-/-^* tissues (versus their respective controls) (**Figure 6A(i,ii)**). Nevertheless, genes up-regulated in *Reverbα^Flox2-6^Adipo^Cre^* and *Reverbα^-/-^* gWAT were significantly enriched around REVERBα ChIP-seq peaks at distances up to 1Mbp (**Figure 6A(i)**), consistent with repression mediated directly by DNA-bound REVERBα. Of these genes, 61.8% were within 100kbp of a stringent REVERBα peak, strongly suggesting that this gene cluster (clock and collagen genes) represents direct targets of REVERBα repression in WAT under NC conditions. Genes up-regulated only in *Reverbα^-/-^* WAT were also enriched around REVERBα peaks (at distances of up to 500kbp) suggesting that at least a proportion of these genes are presumptive direct targets of REVERBα regulation. In contrast, no enrichment of genes with decreased expression in *Reverbα^Flox2-6^Adipo^Cre^* or *Reverbα^-/-^* WAT was evident at any distance from REVERBα peaks (**Figure 6A(ii)**). Thus, REVERBα activation of transcription involves a different mechanism of regulation, likely involving secondary or indirect mechanisms (e.g. de-repression of another repressor), as previously proposed to explain REVERBα transactivation (Le Martelot et al., 2009).

HFD-feeding of *Reverbα^Flox2-6^Adipo^Cre^* mice greatly increased the overlap of differentially-expressed genes with those differentially expressed in *Reverbα^-/-^* WAT. (**Figure 6A(iii,iv)**). We observed a highly significant proximity enrichment of these commonly up-regulated genes (863) to sites of REVERBα chromatin binding (**Figure 6A(iii)**), but again, saw no enrichment of the commonly down-regulated genes (**Figure 6A(iv)**). This integration of transcriptome and cistrome profiling suggests that REVERBα’s exertion of direct repressive control over gene targets is dependent on metabolic state, and increases substantially under HFD-feeding conditions. Of note, the 1714 genes up-regulated only in the *Reverbα^-/-^* WAT retained some proximity enrichment to REVERBα binding sites. This suggests that some of these genes are direct REVERBα targets but subject to additional transcriptional controls.

Consistent with the healthier metabolic phenotype and transcriptome changes observed in obese *Reverbα^Flox2-6^Adipo^Cre^* mice, we found that the large majority of the 863 REVERBα gene targets unmasked by HFD-feeding are normally repressed in obesity (**Figure 6B**), with 551 (63.8%) being significantly down-regulated in obese control (Cre^-ve^) animals. These genes include important regulators of lipid and mitochondrial metabolism (**Figure 6C**) - including *Fasn, Scd1, Dgat2, Cs* - and FGF-21 co-receptor *Klb*, a previously-identified REVERBα target gene (Jager et al., 2016). Considered together, these findings suggest that the healthy adiposity phenotype in HFD-fed *Reverbα^Flox2-6^Adipo^Cre^* mice results from de-repression of REVERBα-controlled pathways allowing continued and efficient lipid synthesis and storage, thus permitting greater expansion of the adipose bed and attenuation of obesity-related dysfunction.

## DISCUSSION

We set out to define the role of REVERBα in the regulation of white adipose tissue metabolism, and subsequently reveal a new understanding of REVERBα function. Together, our data show REVERBα to be a state-dependent regulator of WAT metabolism, with its widespread repressive action only unmasked by diet-induced obesity. Surprisingly, *Reverbα* expression in WAT appears to limits the energy buffering function of the tissue. This finding parallels our recent work in the liver (Hunter et al., 2020). Hepatic-selective loss of REVERBα carries no metabolic consequence, with REVERBα-dependent control over hepatic energy metabolism revealed only upon altered feeding conditions. Contrary to current understanding, our findings therefore suggest that REVERBα (and potentially other components of the peripheral clock) does not impose rhythmic repression of metabolic circuits under basal conditions, but rather determines tissue responses to altered metabolic state.

As reported previously by us and others (Delezie et al., 2012; Hand et al., 2015), global deletion of *Reverbα* leads to an increase in lipogenesis, adipose tissue expansion, and an exaggerated response to diet-induced obesity. How the loss of REVERBα specifically in WAT contributes to this phenotype has not previously been addressed. Here, we use proteomic, transcriptomic and lipid profiling studies to show a clear bias towards fatty acid synthesis and triglyceride storage within *Reverbα^-/-^* WAT. Adipocyte-targeted deletion of *Reverbα* reveals only a modest phenotype, and a relatively selective set of gene targets, limited to clock processes and collagen dynamics. These genes are concurrently de-regulated in *Reverbα^-/-^* adipose, and are found in proximity to REVERBα binding sites, strongly implicating them as direct targets of REVERBα repressive activity.

The reduced inflammation seen in obese *Reverbα^Flox2-6^Adipo^Cre^* mice is likely multifactorial, but may be secondary to a reduction in the pro-inflammatory free fatty acid pool, resulting from de-repression of lipogenic and mitochondrial metabolism pathways, or the absence of signals from dead/dying adipocytes. It is of note that improved metabolic flexibility is proposed to be beneficial in other mouse models of metabolic disease (Jonker et al., 2012; Kim et al., 2007; Virtue et al., 2018) and in human obesity (Aucouturier et al., 2011; Begaye et al., 2020). We identify extracellular matrix as a new direct target of adipose REVERBα action; altered regulation of WAT collagen production and modification is also likely to contribute to the rapid and continued adipose tissue expansion and reduced obesity-related fibrosis. The clock has been linked to ECM re-modelling in other tissues (Chang et al., 2020; Dudek et al., 2015; Sherratt et al., 2019), where it is thought to coordinate ECM dynamics, collagen turnover and secretory processes (Chang et al., 2020). Adipose-specific deletion of *Reverbα* now provides a unique model to explore the complex ECM responses which accompany obesity-related tissue hypertrophy and development of fibrosis.

A role for the clock in the regulation of WAT function has been reported in the literature (Barnea et al., 2015; Paschos et al., 2012; Shostak et al., 2013), perhaps implying that it is the rhythmicity conferred by the clock which is important for WAT metabolism. However, despite robust rhythms of clock genes persisting, rhythmic gene expression in gWAT is largely attenuated following genetic disruption of SCN function (Kolbe et al., 2016). This supports the alternative notion that an intact local clock is not the primary driver of rhythmic peripheral tissue metabolism. Indeed, metabolic processes, including lipid biosynthesis, were highly enriched in the cohort of SCN-dependent rhythmic genes from this study (Kolbe et al., 2016), implying that feeding behaviour and WAT responses to energy flux are more important than locally-generated rhythmicity for adipose function.

The modest impact of adipose-selective *Reverbα* deletion is both at odds with the large effect of global *Reverbα* deletion, and with the extensive WAT cistrome identified for REVERBα (even by stringent peak calling as we have done here). This suggests that the tissue-specific actions of REVERBα are necessary, but not sufficient, and require additional regulation from the metabolic state. Although not explored to the same extent, a lipogenic phenotype of liver-specific RORα/γ deletion has previously been shown to be unmasked by HFD-feeding (Zhang et al., 2017). By driving adipose tissue hypertrophy through HFD-feeding of the *Reverbα^Flox2- 6^Adipo^Cre^* mice, we observed a stark difference in the adipose phenotype of targeted mice and littermate controls. WAT tissue lacking *Reverbα* showed significantly increased tissue expansion, but little evidence of normal obesity-related pathology (tissue fibrosis, and immune cell infiltration/inflammation). Genes controlling mitochondrial activity, lipogenesis and lipid storage were relatively spared from the obesity-related down-regulation observed in control mouse tissue, and were associated with the WAT REVERBα cistrome. Thus, in response to HFD-feeding, REVERBα acts to repress metabolic activity in the adipocyte and limit tissue expansion (albeit at the eventual cost of tissue dysfunction, inflammation and development of adipose fibrosis).

The broadening of REVERBα’s regulatory influence in response to obesity likely reflects a change to the chromatin environment in which REVERBα operates (**Figure 6 Supplementary B**). The majority of emergent REVERBα target genes are repressed in obese adipose when *Reverbα* expression is intact (**Figure 6B**). As these genes are not de-repressed by *Reverbα* loss in normal adipose, REVERBα activity must be redundant or ineffective in a ‘basal’ metabolic state. Subsequent emergence of REVERBα’s transcriptional control may reflect alterations in chromatin accessibility or organisation, and/or the presence of transcriptional repressors and accessory factors required for full activity. As REVERBα is itself proposed to regulate enhancer-promoter loop formation (Kim et al., 2018), modulation of *Reverbα* expression would be a further important variable here. Such reshaping of the regulatory landscape likely occurs across tissues, and may explain why, with metabolic challenge, emergent circadian rhythmicity is observed in gene expression (Eckel-Mahan et al., 2013; Kinouchi et al., 2018; Tognini et al., 2017), and in circulating and tissue metabolites (Dyar et al., 2018; Eckel-Mahan et al., 2013).

Here, we now uncover a role for REVERBα in limiting the energy-buffering role of WAT, a discovery which may present therapeutic opportunity as we cope with an epidemic of human obesity. Despite recent findings which have cast doubt on the utility of some of the small molecule REVERBα ligands (Dierickx et al., 2019), antagonising WAT REVERBα now emerges as a potential target in metabolic disease. Finally, our study suggests that a functioning circadian clock may be beneficial in coping with acute mistimed metabolic cues but, that under chronic energy excess, may contribute to metabolic dysfunction and obesity- related pathology.

## MATERIALS AND METHODS

### Animal experiments

All experiments described here were conducted in accordance with local requirements and licenced under the UK Animals (Scientific Procedures) Act 1986, project licence number 70/8558 (DAB). Procedures were approved by the University of Manchester Animal Welfare and Ethical Review Body (AWERB). Unless otherwise specified, all animals had *ad libitum* access to standard laboratory chow and water, and were group-housed on 12hour:12hour light:dark (LD) cycles and ambient temperature of 22°C+/-1.5°C. Male mice (*Mus musculus*) were used for all experimental procedures. All proteomics studies were carried out on 13-week-old weight matched males. RNA-seq studies for Figure 3 were carried out on 12-14 week-old weight-matched males; the RNA-seq study for Figure 5 was carried out on 28 week-old males (following 16 weeks of high-fat diet or normal chow feeding).

### Reverbα^-/-^

*Reverbα^-/-^* mice were originally generated by Ueli Schibler (University of Geneva) (Preitner et al., 2002). These mice were created by replacing exons 2-5 of the *Reverbα* gene by an in-frame LacZ allele. Mice were then imported to the University of Manchester and backcrossed to C57BL/6J mice.

### _Reverbα_Flox2-6

A CRISPR-Cas9 approach was used to generate a conditional knock allele for *Nr1d1* (*Reverbα*), as described (Hunter et al., 2020). LoxP sites were integrated, in a two-step process, at intron 2 and intron 6, taking care to avoid any previously described transcriptional regulatory sites (Yamamoto et al., 2004). A founder animal with successful integration of both the 5’ and 3’ loxP sites, transmitting to the germline, was identified and bred forward to establish a colony.

### Reverbα^Flox2-6^Adipo^Cre^

*Adiponectin*-driven Cre-recombinase mice (Eguchi et al., 2011; Jeffery et al., 2014) were purchased from The Jackson Laboratory and subsequently bred against the *Reverbα^Flox2-6^* at the University of Manchester.

### In vivo phenotyping

Body composition of mice was analysed prior to cull by quantitative magnetic resonance (EchoMRI 900). Energy expenditure was measured via indirect calorimetry using CLAMS (Columbus Instruments) for 10-12 week old male mice. Mice were allowed to acclimatise to the cages for two days, prior to an average of 5 days of recordings being collected. Recording of body temperature and activity was carried out via surgically implanted radiotelemetry devices (TA-F10, Data Sciences International). Data is shown as a representative day average of single- housed age-matched males. For the diet challenge, male mice were fed high-fat diet (HFD; 60% energy from fat; DIO Rodent Purified Diet, IPS Ltd) for a period of 10-16 weeks from 12 weeks of age. Blood glucose was measured from tail blood using the Aviva Accuchek meter (Roche). For the insulin tolerance test, mice were fasted from ZT0, then injected with 0.75 IU/kg human recombinant insulin (I2643, Sigma Aldrich) at ZT6 (time “0 minutes”).

### Insulin ELISA

Insulin concentrations were measured by ELISA (EZRMI-13K Rat/Mouse insulin ELISA, Merck Millipore) according to the manufacturer’s instructions. Samples were diluted in matrix solution to fall within the range of the assay. Internal controls supplied with the kit were run alongside the samples and were in the expected range.

### Histology

gWAT was collected and immediately fixed in 4% paraformaldehyde for 24 hours, transferred into 70% ethanol, and processed using a Leica ASP300 S tissue processor. 5μm sections underwent H&E staining (Alcoholic Eosin Y solution (HT110116) and Harris Haematoxylin solution (HHS16), Sigma Aldrich), Picrosirius Red staining (see below), or F4/80 immunohistochemistry (see below), prior to imaging using a Snapshots Olympus single slide scanner at 10x or 20x objective magnification alongside Olympus cellSens Dimension software (version 1.18). Percentage area stained was quantified using ImageJ (version 1.52a) as detailed in the online ImageJ documentation (https://imagej.nih.gov/ij/docs/examples/stained-sections/index.html), with 5-12 images quantified per animal. Adipocyte area was quantified using the Adiposoft ImageJ plug-in (version 1.16 - https://imagej.net/Adiposoft).

For Picrosirius Red staining, sections were dewaxed and rehydrated using the Leica ST5010 Autostainer XL. Sections were washed in distilled water and then transferred to Picrosirius Red (Direct Red 80, Sigma Aldrich) (without the counterstain) for 1 hour. Sections were then washed briefly in 1% acetic acid. Sections were then dehydrated, cleared and mounted using the Leica ST5010 Autostainer XL.

For F4/80 immunohistochemistry, sections were dewaxed and rehydrated prior to enzymatic antigen retrieval (trypsin from porcine pancreas (T7168, Sigma)). Sections were treated with 3% hydrogen peroxide to block endogenous peroxidase activity followed by further blocking with 5% goat serum. Rat mAb to F4/80 (1:500) (ab6640, Abcam) was added and sections were incubated overnight at 4°C. Sections were washed before addition of the biotinylated anti-rat IgG (BA-9400, H&L) secondary antibody (1:1500) for 1 hour. Sections were developed using VECTAstain® Elite® ABC kit peroxidase, followed by DAB Peroxidase substrate (Vector Labs) and counterstained with haematoxylin. Slides were then dehydrated, cleared and mounted.

### Lipid extraction and gas chromatography

Total lipid was extracted from tissue lysates using chloroform-methanol (2:1; v/v) according to the Folch method (Folch et al., 1957). An internal standard (tripentadecanoin glycerol (15:0)) of known concentration of was added to samples for quantification of total triacylglyceride. Lipid fractions were separated by solid-phase extraction and fatty acid methyl esters (FAMEs) were prepared as previously described (Heath et al., 2003). Separation and detection of total triglyceride FAMEs was achieved using a 6890N Network GC System (Agilent Technologies; CA, USA) with flame ionization detection. FAMEs were identified by their retention times compared to a standard containing 31 known fatty acids and quantified in micromolar from the peak area based on their molecular weight. The micromolar quantities were then totalled and each fatty acid was expressed as a percentage of this value (molar percentage; mol%).

### Proteomics

Mice were culled by cervical dislocation and the gWAT was immediately removed and washed twice in ice-cold PBS and then once in ice-cold 0.25M sucrose, prior to samples being snap-frozen in liquid nitrogen and stored at -80°C. To extract the protein, the samples were briefly defrosted on ice and then cut into 50mg pieces and washed again in ice-cold PBS. The sample was then lysed in 200μl of 1M Triethylammonium bicarbonate buffer (TEAB; Sigma) with 0.1% (w/v) sodium dodecyl sulphate (SDS) with a Tissue Ruptor (Qiagen). Samples were centrifuged for 5 minutes, full speed, at 4°C and the supernatant collected into a clean tube. A Biorad protein assay (Biorad) was used to quantify the protein and Coomassie protein stain (InstantBlue™ Protein Stain Instant Blue, Expedeon) to check the quality of extraction. Full methods of subsequent iTRAQ proteomic analysis including bioinformatic analysis has been published previously (Xu et al., 2019). Here the raw data was searched against the mouse Swissprot database (release October 2017) using the paragon algorithm on Protein-Pilot (Version 5.0.1, AB SCIEX). A total of 33847 proteins were searched. As described (Xu et al., 2019), Bayesian protein-level differential quantification was performed by Andrew Dowsey (University of Bristol) using their own BayesProt (version 1.0.0), with default choice of priors and MCMC settings. Expression fold change relative to the control groups were determined and proteins with a global false discovery rate of >0.05 were deemed significant.

### 3T3-L1 cells

3T3-L1 cells (ATCC) were maintained in Dulbecco’s Modified Eagle’s Medium - high glucose (DMEM/D6429, Sigma-Aldrich) supplemented with 10% Foetal Bovine Serum and 1% penicillin/streptomycin at 37°C/5% CO2. Cells were grown until confluent, passaged and plated into 12-well tissue culture plates for differentiation. The differentiation protocol was initiated 5 days later. Cells were treated with 10µg/mL insulin (Sigma-Aldrich), 1µM dexamethasone (Sigma-Aldrich), 1µM rosiglitazone (AdooQ Bioscience) and 0.5mM IBMX (Sigma-Aldrich) prepared in DMEM + 10% FBS + 1% Pen/Strep for 3 days. On day 3 and day 5 the cell culture media was changed to 10µg/mL insulin and 1µM rosiglitazone in DMEM + 10% FBS + 1% Pen/Strep. On day 7, the cell culture media was changed to 10µg/mL insulin in DMEM + 10% FBS + 1% Pen/Strep. Finally on day 10, the cell culture media was changed to DMEM + 10% FBS + 1% Pen/Strep without any additional differentiation mediators. Cells were used from day 11 onwards. Lipid droplets were visible by day 5.

For knockdown studies, mature 3T3-L1 adipocytes were transfected with Sicontrol (Control ON-TARGETplus siRNA, Dharmacon), SiReverbα (Mouse NR1D1 ON-TARGETplus siRNA, Dharmacon) or SiReverbβ (Mouse NR1D2 ON-TARGETplus siRNA, Dharmacon) at 50nM concentration using Lipofectamine RNAiMAX (Invitrogen) as a transfection reagent. Briefly, 12-well plates were coated with poly-L-lysine hydrobromide (Sigma) and incubated for 20-30 minutes prior to excess poly-L-lysine being removed and the plates allowed to dry. SiRNAs and RNAiMAX transfection reagent were separately mixed with reduced serum media (Opti-MEM, Gibco). The control or Reverbα/β SiRNA was then added to each well and mixed with an equal quantity of RNAiMAX and then incubated for 5 minutes at room temperature. Mature 3T3-L1 adipocytes were trypsinised (trypsin-EDTA solution, Sigma) and resuspended in FBS without P/S prior to being re-plated into the wells containing the SiRNA. After 24 hours the transfection mix was removed and replaced with DMEM without FBS or P/S. The cells were then collected 48 hours after transfection.

### RNA extraction (cells)

RNA was extracted from cells using the ReliaPrep™ RNA Cell Miniprep system (Promega, UK), following manufacturer’s instructions. RNA concentration and quality was determined with the use of a NanoDrop spectrophotometer and then stored at -80°C.

### RNA extraction (tissue)

Frozen adipose tissue was homogenised in TRIzol Reagent (Invitrogen) using Lysing Matrix D tubes (MP Biomedicals) and total RNA extracted according to the manufacturer’s TRIzol protocol. To remove excess lipid, samples then underwent an additional centrifugation (full speed, 5 minutes, room temperature) prior to chloroform addition. For the RNA sequencing samples the isopropanol phase of TRIzol extraction was transferred to Reliaprep tissue Miniprep kit (Promega, USA) columns to ensure high quality RNA samples were used. The column was then washed, DNAse treated and RNA eluted as per protocol. RNA concentration and quality was determined with the use of a NanoDrop spectrophotometer and then stored at -80°C. For RNA-seq, RNA was diluted to 1000ng in nuclease-free water to a final volume of 20uL.

### RT-qPCR

For RT-qPCR, samples were DNase treated (RQ1 RNase-Free DNase, Promega, USA) prior to cDNA conversion High Capacity RNA-to-cDNA kit (Applied Biosystems). qPCR was performed using a GoTaq qPCR Master Mix (Promega, USA) and primers listed in Supplementary Table 1 using a Step One Plus (Applied Biosystems) qPCR machine. Relative quantities of gene expression were determined using the [delta][delta] Ct method and normalised with the use of a geometric mean of the housekeeping genes *Hprt, Ppib* and *Actb*. The fold difference of expression was calculated relative to the values of control groups.

### RNA-seq

Adipose tissue was collected from adult male mice (n=6-8 per group) at ZT8 and flash-frozen. Total RNA was extracted and DNase-treated as described above. Biological replicates were taken forward individually to library preparation and sequencing. For library preparation, total RNA was submitted to the Genomic Technologies Core Facility (GTCF). Quality and integrity of the RNA samples were assessed using a 2200 TapeStation (Agilent Technologies) and then libraries generated using the TruSeq® Stranded mRNA assay (Illumina, Inc.) according to the manufacturer’s protocol. Briefly, total RNA (0.1-4μg) was used as input material from which polyadenylated mRNA was purified using poly-T, oligo-attached, magnetic beads. The mRNA was then fragmented using divalent cations under elevated temperature and then reverse transcribed into first strand cDNA using random primers. Second strand cDNA was then synthesised using DNA Polymerase I and RNase H. Following a single ’A’ base addition, adapters were ligated to the cDNA fragments, and the products then purified and enriched by PCR to create the final cDNA library. Adapter indices were used to multiplex libraries, which were pooled prior to cluster generation using a cBot instrument. The loaded flow-cell was then paired-end sequenced (76 + 76 cycles, plus indices) on an Illumina HiSeq4000 instrument. Finally, the output data was demultiplexed (allowing one mismatch) and BCL-to-Fastq conversion performed using Illumina’s bcl2fastq software, version 2.17.1.14

### RNA-seq data processing & differential gene expression analysis

Paired-end RNA-seq reads were quality assessed using FastQC (v 0.11.3), FastQ Screen (v 0.9.2) (Wingett and Andrews, 2018). Reads were processed with Trimmomatic (v 0.36) (Bolger et al., 2014) to remove any remaining sequencing adapters and poor quality bases. RNA-seq reads were then mapped against the reference genome (mm10) using STAR (version 2.5.3a) (Dobin et al., 2013). Counts per gene (exons) were calculated by STAR using the genome annotation from GENCODEM16. Differential expression analysis was then performed with edgeR (Robinson et al., 2010) using QLF-tests based on published code (Chen et al., 2016). Changes were considered significant if they reached a FDR cut-off of <0.05. Interaction analysis was performed with stageR (Van den Berge et al., 2017) in conjunction with Limma voom (Law et al., 2014), setting alpha at 0.05.

### Published ChIP-seq

Raw gWAT ChIP-seq data (Zhang et al., 2015) was downloaded from the GEO Sequence Read Archive (GSE67973) using the sratoolkit package (v2.9.2) (*fastq-dump* tool). The following datasets were used: REVERBα ZT10 ChIP-seq (two replicates) - SRR1977510, SRR1977511 – and ZT10 input raw data - SRR1977512. Reads were aligned to the mm10 genome with Bowtie2 (v.2.3.4.3) (Langmead and Salzberg, 2012), then sorted, indexed BAM files were produced with SAMtools (v.1.9) (Li et al., 2009). MACS2 (Zhang et al., 2008) was used to call peaks from the experimental BAM files (-t) against the input control BAM file (-c), with the following options specified: *-f BAMPE -g mm --keep-dup=1 -q 0.01 -- bdg --SPMR --verbose 0*. High-stringency peaks were defined as those with >7.5 fold-enrichment over input.

### Integrating RNA-seq and ChIP-seq

In order to calculate enrichment of RNA-seq-based gene clusters with respect to ChIP-seq peaks, we used our in-house custom tool (Yang et al., 2019) which calculates gene cluster enrichment within specified distances from the centre of peaks (see also *Code Availability* statement below). The tool extends peaks in both directions for the given distances and extracts all genes whose TSSs overlap with the extended peaks. Given these genes, the inputted RNA-seq-based gene cluster, and the overlap of these two groups, it performs a hypergeometric test with the total number of genes in the mm10 genomes as background.

### Pathway analysis

Pathway enrichment analysis of ENTREZ gene identifiers, either extracted from RNA-seq or proteomics data, was carried out using the R Bioconductor package *ReactomePA* (Yu and He, 2016). The *enrichPathway* tool was used with the following parameters: *organism = “mouse”, pAdjustMethod = “BH”, maxGSSize = 2000, readable = FALSE.* We considered pathways with a Padj<0.01 to be significantly enriched. For Figure S3A, no pathways had Padj<0.01 (likely due to the small number of inputted genes), hence we show the 5 pathways with the smallest Padj values.

### Protein extraction and Western blotting

Small pieces (<100mg) of tissue were homogenised with the FastPrep Lysing Matrix D system (MP Biomedicals) in T-PER (Thermo Fisher Scientific), supplemented with protease inhibitor cocktail (Promega) at 1:50 dilution. Benzonase nuclease (EMD Millipore) was added (2μl), the homogenate briefly vortexed, then incubated on ice for 10 minutes. Homogenates were then centrifuged for 8 minutes at 10,000g, at 4°C, and the supernatant removed (avoiding any lipid layer). Protein concentration was quantified using the Bio-Rad Protein Assay (Bio-Rad). For Western blotting, equal quantities (75μg for detection of REVERBα) of protein were added to 4x NuPAGE LDS sample buffer (Invitrogen), NuPAGE sample reducing agent (dithiothreitol) (Invitrogen) and water, and denatured at 70°C for 10 minutes. Samples were run on 4-20% Mini-PROTEAN TGX Precast Protein Gels (Bio-Rad) before wet transfer to nitrocellulose membranes. Membranes were blocked with Protein-Free Blot Blocking Buffer (Azure Biosystems), and subsequent incubation and wash steps carried out following manufacturer’s instructions. Primary and secondary antibodies used were as listed in Supplementary Table 2, with primary antibodies being used at a 1:1000 dilution and secondary antibodies at 1:10,000. Membranes were imaged using chemiluminescence or the LI-COR Odyssey system. Uncropped blot images can be provided by the corresponding author upon request.

### Statistics

To compare two or more groups, t-tests or ANOVAs were carried out using GraphPad Prism (v.8.4.0). For all of these, the exact statistical test used and n numbers are indicated in the figure legends. All n numbers refer to individual biological replicates (ie. individual animals). Unless otherwise specified, bar height is at mean, with error bars indicating +/-SEM. In these tests, significance is defined as *P<0.05 or **P<0.01 (P values below 0.01 were not categorised separately, i.e. no more than two stars were used, as we deemed this to be a meaningful significance cut-off). Statistical analyses of proteomics, RNA-seq and ChIP-seq data were carried out as described above in Methods, using the significance cut-offs mentioned. Plots were produced using GraphPad Prism or R package ggplot2.

### Data availability

RNA-seq data generated in the course of this study has been uploaded to ArrayExpress and is available at http://www.ebi.ac.uk/arrayexpress/experiments/E-MTAB-1. 8840. Raw proteomics data has been uploaded to Mendeley Data (https://data.mendeley.com/datasets/wskyz3rhsg/draft?a=ef40a1ec-36a4-4509-979d-32d494b96585). Other data supporting the findings of this study are available from the corresponding author upon reasonable request.

### Code availability

The custom Python code (Yang et al., 2019) used to carry out the peaks- genes enrichment analysis in this study is available at http://bartzabel.ls.manchester.ac.uk//Pete/PF5HZns0zv/pegs-0.2.0.tgz.

## Acknowledgements

We thank Rachel Scholey, I-Hsuan Lin, Ping Wang and Peter Briggs (Bioinformatics Core Facility, UoM), and Thea Danby (Faculty of Biology, Medicine and Health, UoM) for statistical and technical assistance, and acknowledge support of core facilities at the University of Manchester: Genomic Technologies Core Facility, Biological Services Unit, and Histological Services Unit. We also acknowledge and thank the support of our funders: the BBSRC (BB/I018654/1 to D.A.B.), the MRC (Clinical Research Training Fellowship MR/N021479/1 to A.L.H.; MR/P00279X/1 to D.A.B; MR/P011853/1 and MR/P023576/1 to D.W.R.), and the Wellcome Trust (107849/Z/15/Z, 107851/Z/15/Z).

## Author Contributions

Conceptualisation, A.L.H., C.E.P., D.W.R., D.A.B.; Methodology, A.A., R.D.U., D.W.R., D.A.B.; Software, M.I., A.L.H.; Investigation, A.L.H., C.E.P., P.D., T.C., P.S.C., N.J.B., R.C.N.; Formal Analysis, A.L.H., C.E.P., R.D.U., L.H., M.I., D.A.B.; Writing, A.L.H., C.E.P., A.S.I.L., D.W.R., D.A.B.; Funding Acquisition, A.L.H., D.W.R., D.A.B.; Supervision, D.A.B.

---

**SUPPLEMENTARY TABLE 1.**
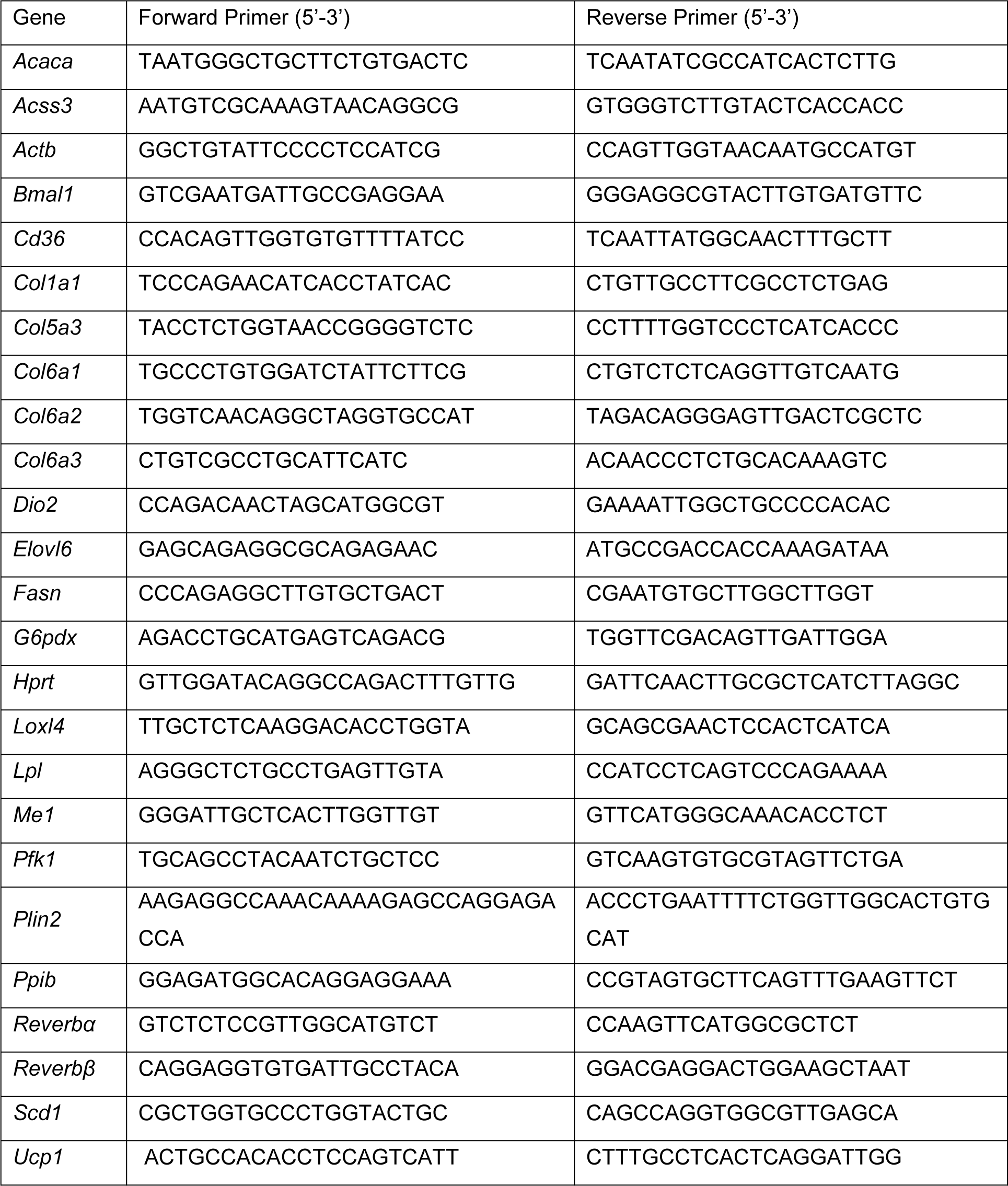
qPCR primer sequences.

**SUPPLEMENTARY TABLE 2.** Antibodies.

| Source and target | Manufacturer | Catalogue identifier, lot | Use |
| --- | --- | --- | --- |
| Rabbit polyclonal anti-REVERBα (NR1D1) | ProteinTech | Cat#14506-1-AP, lot 5745;<br>RRID:AB_11182604 | Western blot, primary |
| Mouse monoclonal anti-UCP1 (clone 536435) | R&D Systems | Cat#MAB6158, lot CCNV0218091;<br>RRID:AB_10572490 | Western blot, primary |
| Mouse monoclonal anti-β-ACTIN (ACTB) | ProteinTech | Cat#60008-1, lot 0001084;<br>RRID:AB_2289225 | Western blot, primary |
| Sheep Anti-Mouse IgG, HRP-linked whole Ab | GE Healthcare | Cat#NA931V, lot 16908225;<br>RRID:AB_772210 | Western blot, secondary |
| Donkey Anti-Rabbit IgG, HRP-linked whole Ab | GE Healthcare | Cat#NA934V, lot 16921443;<br>RRID:AB_772206 | Western blot, secondary |
| Goat polyclonal Anti-Rabbit IgG (H+L), DyLight 680 Conjugate | Cell Signaling Technology | Cat#5366P, lot 7;<br>RRID:AB_10693812 | Western blot, secondary |
| Rat monoclonal anti-F4/80 | Abcam | Cat#ab6640, lot 845724;<br>RRID:AB_1140040 | Immunohistochemistry, primary |
| Goat biotinylated anti-rat IgG (affinity purified) | H&L | Cat#BA-9400, lot ZB1216;<br>RRID:AB_2336202 | Immunohistochemistry, secondary |

